# Artificial selection of stable rhizosphere microbiota leads to heritable plant phenotype changes

**DOI:** 10.1101/2021.04.13.439601

**Authors:** Samuel Jacquiod, Aymé Spor, Shaodong Wei, Victoria Munkager, David Bru, Søren J. Sørensen, Christophe Salon, Laurent Philippot, Manuel Blouin

## Abstract

Research on artificial selection of microbial community has become popular due to perspectives in improving plant and animal health^1-4^. However, reported results still lack consistency^5-8^. We hypothesized that artificial selection may provide desired outcomes provided that microbial community structure has stabilized along the selection process. In a ten-generation artificial selection experiment involving 1,800 plants, we selected rhizosphere microbiota of *Brachypodium distachyon* that were associated with high or low levels of leaf greenness, a proxy for plant health^9^. Monitoring of the rhizosphere microbiota dynamics showed strong oscillations in community structure during an initial transitory phase of five generations, with no heritability in the selected property. In the last five generations, the structure of microbial communities displayed signs of stabilization, concomitantly to the appearance of heritability in leaf greenness. Selection pressure, initially ineffective, became successful in changing the greenness index in the intended direction, especially toward high greenness values. We showed a remarkable congruence between plant traits and selected microbial community structures, highlighting two phylogenetically distinct microbial sub-communities correlating with leaf greenness, whose abundance was significantly steered by directional artificial selection. Understanding microbial community structure stabilization can thus help improve the reliability of artificial microbiota selection.

**Short Sentence:** Stable microbiota selection enables trait heritability

## Main Text

Empirical studies of artificial selection have demonstrated that it is possible to steer microbiota across generations to modify microbial ecosystems properties^4-8^. In some cases, the selected property can be a trait displayed by the host of a microbiota, like with microbial communities associated to plants^5,10-13^. This opens new avenues for plant breeding *via* directional artificial selection of rhizosphere microbiota^1-4^. Still, while previous studies reported significant selection effects^6,7,11^, the selected property may not be perennial and lost during the process^6,9^. From a practical point of view, it is thus crucial to understand the causes behind this inconsistency. We hypothesized that an important prerequisite for successful selections is to reach a stable state in microbial community structure^14^. In the field of artificial selection of communities^15-17^, mathematical models have shown that the heritability of the selected property, one of the three essential features of a unit of selection^18^, depends on the stability of community structure^19,20^. However, this hypothesis about ‘stability of community structure being a prerequisite for the heritability of the selected property’ has never been empirically confirmed. Here, we tested it by investigating the dynamics of microbial community and the heritability in the selected property during the selection process.

We artificially selected rhizosphere microbiota according to their impact on a leaf greenness index, a remote proxy for plant nutritional and health status^9^ (Fig. S1), using the grass species *Brachypodium distachyon* grown in microcosms in a climatic chamber (Fig. S2-S3). There were two treatments, in which low and high levels of leaf greenness were selected for, respectively (hereafter, the low and high selection groups). There was also a control group, in which selection was random. The selection process was repeated across ten generations, each lasting four weeks (Fig. S2). The three experimental groups (low selection, high selection, and control) each contained three independent replicate lineages composed of twenty microcosms. At the end of each generation, 3 of the 20 microcosms within each lineage were selected based on their leaf greenness values (Fig. S4). Their rhizosphere microbiota were extracted, pooled, and used to inoculate seedlings of the next generation (Fig. S2-S3). We used the same seed batch throughout the experiment, thus only the rhizosphere microbiota could evolve, not the plant genotype.

Analysis of leaf greenness revealed a generation effect (39.77% of the variance, *P* < 0.001; Table S1, Fig.1, A-B) due to uncontrolled biotic and/or abiotic variations, as commonly observed in this kind of selection experiments^5,12^. Nevertheless, we detected significant changes in leaf greenness due to the microbiota-based artificial selection (selection + lineage = 10.15% of the variance, *P* < 0.001, Tab.S1), occurring at specific generations (Fig. 1, A-B). Across the entire experiment, this resulted in a significant increase in leaf greenness in the high selection group compared to the control, but not for the low selection group (Fig. 1, C). This trend was also observed on the other plant traits acquired by the image analysis (Fig. S4), again with a significant increase only for the high selection group (Fig. S5). The effect of selection on the targeted plant trait was more pronounced once data was standardized with the random selection group to control the generation effect (z-score normalization, Fig. 1, D-E).

**Figure 1:**
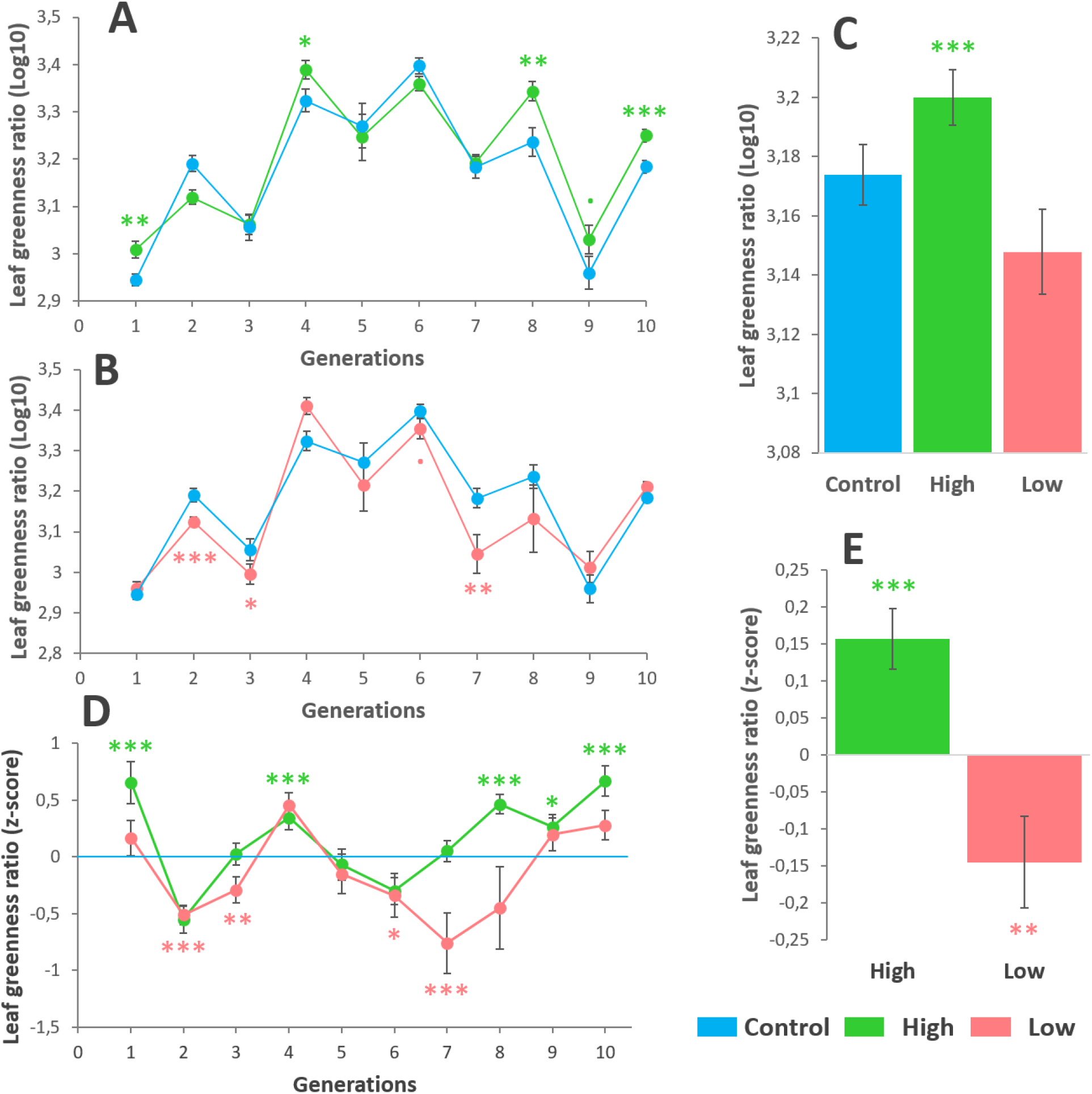
Analysis of the leaf greenness during the course of selection. Panel A and B are showing the evolution of shoot greenness across the ten generations in the high and low selection groups compared to the control group, respectively. Panel C shows the overall averaged leaf greenness values in all selection groups. Panel D shows the evolution of the standardized leaf greenness (z-score using the control group average and standard deviation) for the high and low selection groups across the ten generations. Panel E is showing the overall averaged leaf greenness values in all selection groups for the z-score standardized data. Statistical comparisons were done against the control group with a one-sided, two samples Student test (panels A-C) and a one-sided, one samples Student test for the standardized data (z-score, tested against zero, panels D-E). *P*-value significance: « *** » for *P* < 0.001; « ** » for *P* < 0.01; « * » for *P* < 0.05; «. » for *P* < 0.1. For panel A, B and D: N = 55-60 replicates per group per time point. For panel C end E: N = 591-596 replicates per group. Error-bar are representing the standard error of the mean.

In parallel, our findings underscore strong oscillations of bacterial community structure for the first five generations of selection (dotted line, *R*^*2*^ = 0.56, *P* < 9.99×10^−5^, Fig. 2, A), occurring for all selection groups (Fig. S6). These oscillations were not observed any more from generation G05, denoting a stabilization of bacterial community structure. Fungal community structure abruptly shifted early on and then continued to change at a slower pace for all selection groups (dotted line, *R*^*2*^ = 0.59, *P* < 9.99×10^−5^, Fig. 2, B, Fig. S6). We found that this stabilization was more pronounced for bacteria in the high selection group (Fig. S6), where the effect of selection on the selected property was the most significant (Fig. 1, C). The overall stabilization of the microbiota was also observed on the alpha diversity for both bacteria (Fig. S7) and fungi (Fig. S8), displaying an initial increase and then plateaued in all treatment groups. Since molecular quantification of bacterial and fungal markers were stable across the experiment (Fig. S9), the most parsimonious interpretation of these results is that our iterative microbiota inoculation procedure in all selection groups has decreased the dominance of microbial species initially present, and promoted rare species, resulting in more even communities. To detect an eventual tipping point in community structure over the course of generations, we applied an unsupervised segmented regression analysis on the beta diversity dynamics of each lineage (Fig. S10, A). Results confirmed the presence of a breaking point at generation G05 on average for both bacterial (range: 3.00-6.65; Fig. S11) and fungal (range: 4.00-7.43, Fig. S12) lineages. When considering all data points regardless of lineages, we confirmed that microbial community dissimilarity sharply decreased from generation G01 to G05 and then stabilized (the slope not significantly different from 0 for bacteria and a weaker slope for fungi; Fig. 3, A, Fig. S10, B). This stabilization was not due to a homogenization of microbial communities amongst lineages due to cross or environmental contaminations, as shown by the distinct microbial community structures obtained in each lineage for all selection groups, except the random lineages of bacteria (*P* < 0.05, Fig. S13). These results suggested the existence of two distinct phases during the course of microbiota selection: a transitory phase before generation G05 and thereafter a stabilization phase (Fig. 3, A). The presence of these two phases was also visible on the selected property (Fig. S1, C), and became blatant when analyzing the property for each phase separately, as the leaf greenness index clearly increased significantly in the high selection group after generation G05 (Fig. S14, A), for all three high lineages (Fig. S14, B). Results in the low selection were more contrasted. Pronounced leaf discoloration was observed in lineage LL1 (Fig. S14, B-D), responsible of the overall trend observed in the low selection group (Fig. 1, E; Fig. S14 A). Lineage LL2 did not differ from the control group, while LL3 displayed the opposite trend compared to our expectations (Fig. S14, B), probably due to the lack of bacterial community stabilization (Fig. S11).

**Figure 2:**
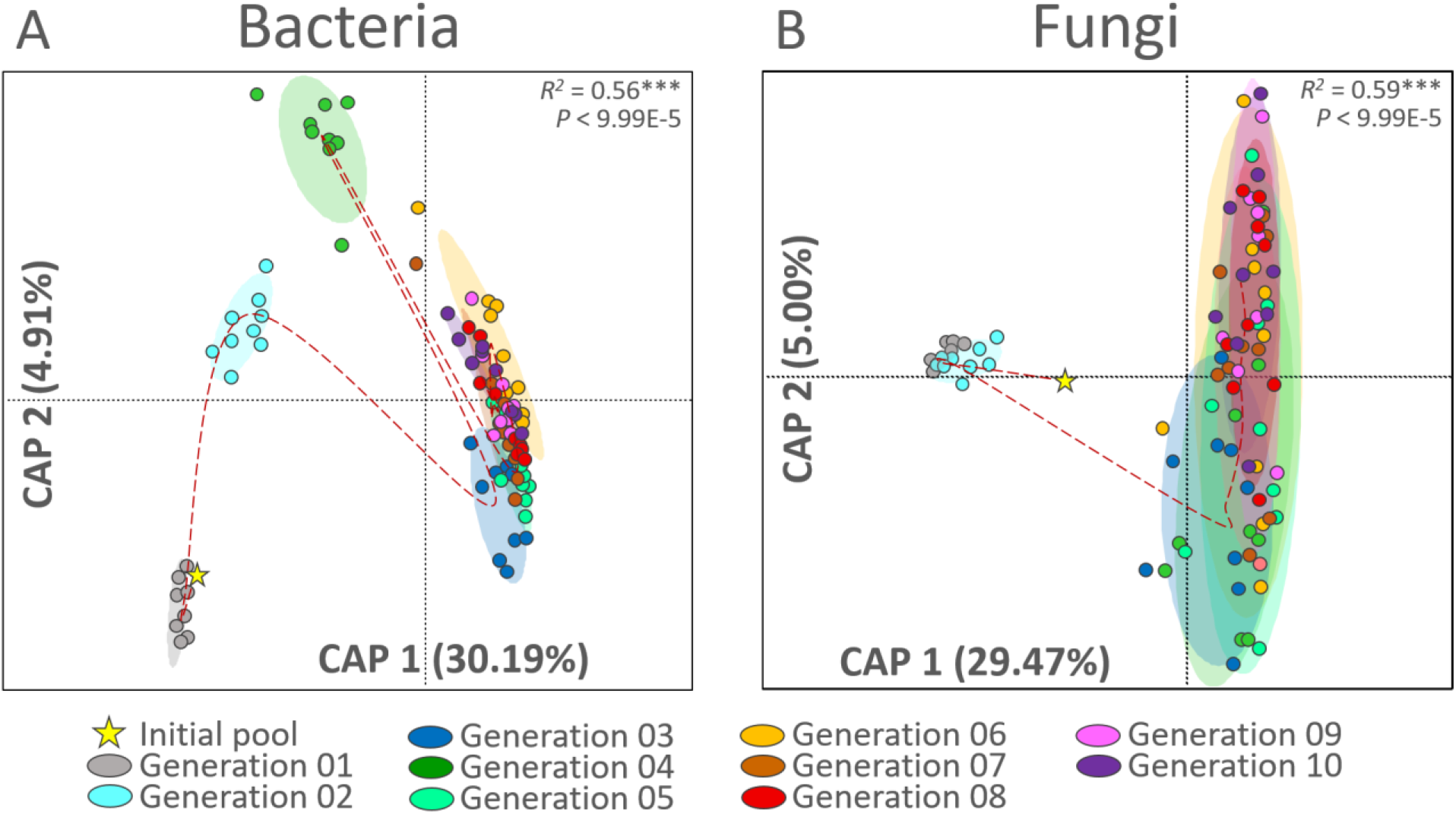
Distance-based redundancy analysis of the microbiota rhizosphere during the course of selection. Panel A and B represent the evolution of community dissimilarity for bacteria and fungi respectively. The same analysis applied for each selection group is available in supporting data (Fig. S6). The models were built using the Bray-Curtis dissimilarity index, with 10.000 group permutations (Bray-Curtis ∼ generation/selection/lineage). The *R*^2^ values are indicating the percentage of variance explained by the model. If significant, the constrained coordinates are shown (model *P* < 0.05, CAP, Constrained Analysis of Principal coordinates). If not, the unsupervised coordinates are shown (model *P* > 0.05, MDS: Multi-Dimensional Scaling).

**Figure 3:**
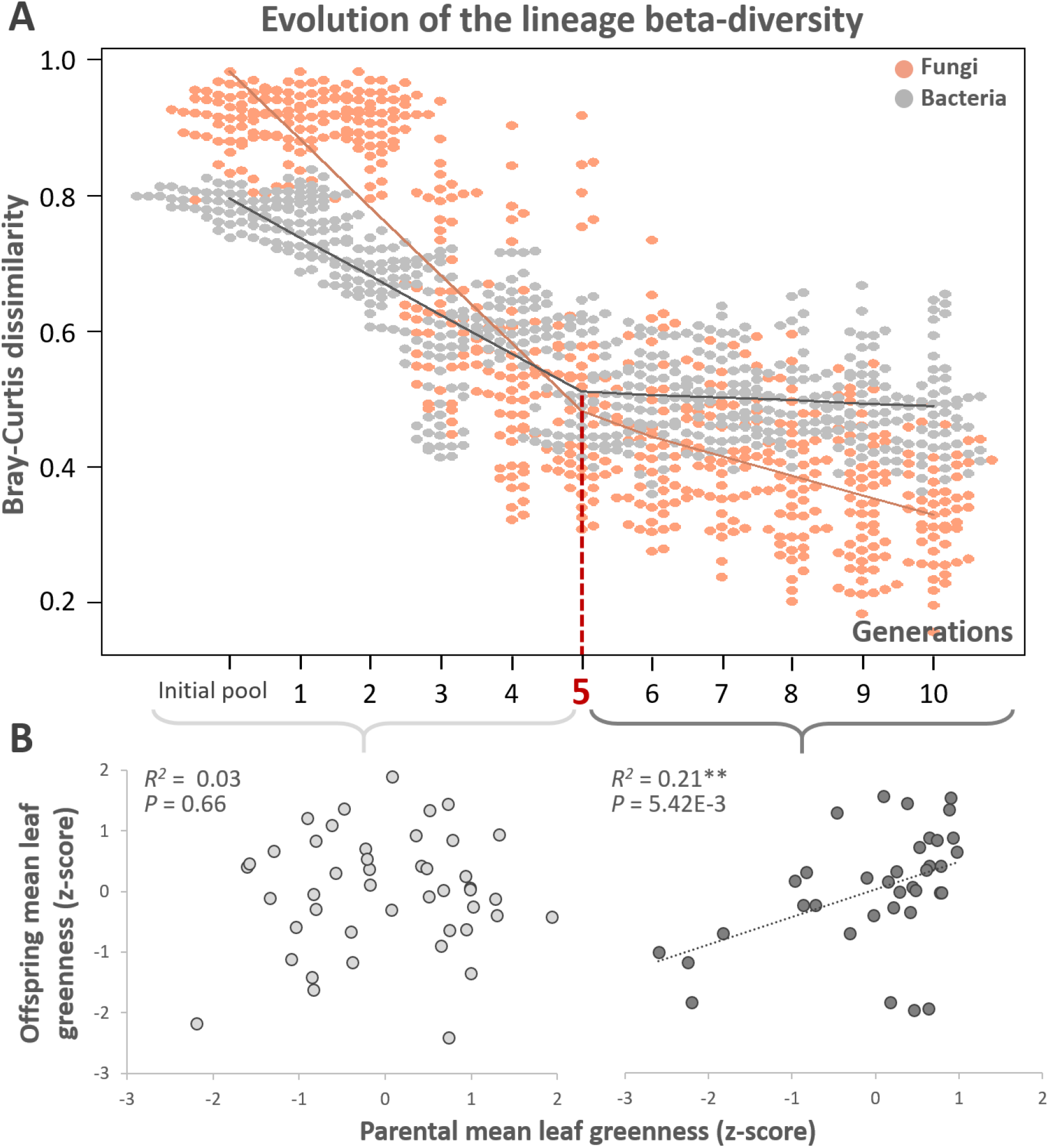
Evolution of the microbial beta diversity and the trait heritability. Panel A shows the overall evolution of each lineage (colored here by bacterial and fungal microbial groups for clarity sake) during the course of selection. The evolution of each bacterial and fungal lineages are displayed in supporting data (Fig. S11-S12). To generate this analysis, we compared six offspring rhizosphere microbiota from G10 in each lineage to their respective pools in descending order until reaching the initial pool used to inoculate the experiment at the beginning (see Fig. S10, A). An unsupervised segmented analysis was performed on each lineage, revealing an average breaking point in the beta diversity slope occurring at generation G05 (Fig. S11-S12). Panel B shows the concomitant evolution of the leaf greenness heritability, calculated as the slope between the averaged selected ‘parent’ phenotype at generation « n » and their averaged ‘offspring’ phenotype at generation « n+1 » in all high, low and control lineages, respectively. A first model was constructed at the transitory phase ([G01-G05], light gray) and a second one at the stabilization phase ([G06-G10], dark gray) according to the beta diversity breaking point. To accurately estimate heritability in our experiment, we integrated values from all lineages (high, low and control) during the [G01-G05] and [G06-G10] intervals based on our unsupervised segmented analysis to spawn sufficient variability to be able to detect whether or not a relationship existed between selected parents and offspring plants. The linear equation for the stabilization phase was y = 0.454x - 6E-16.

Our hypothesis was that stability in microbial community structure is a prerequisite to the heritability of the selected property. Here, we considered community heritability as the regression coefficient between the parental and offspring values of the selected property^7^. We calculated this regression across all selection groups to get both low, random and high values of parent/offspring couples, for the transitory and stabilization phases respectively (Fig. 3, B). According to our prediction, we observed no significant correlations between greenness indices of the parental and offspring microbial communities during the transitory phase (*R*^2^ = 0.03, *P* = 0.66), but a significant correlation during the stabilization phase (*R*^2^ = 0.21, *P* =5.42×10^−3^). We also noted that community dissimilarity in the high selection lineages, for which selection was efficient, was significantly lower compared to the other groups (Fig. S10, C). Taken together, these results confirm modeling predictions that successful artificial selection on microbial communities requires stability, thus enabling a heritable property^19,20^. Indeed, during the transitory phase, as each microbial species had its own population dynamic depending on biotic and abiotic factors, there was no *a priori* synchronicity between their abundance variations. Directional artificial selection applied during this phase resulted in the selection of microbial community that i) may contribute to the observed plant property and ii) reached a certain degree of synchronicity amongst the multiple population dynamics, thus leading to the emergence of a reproduced pattern in microbial community structures that will ensure the heritability of the selected property across generations^19,20^. Microbial species not showing synchronicity might be selected once, but not over the course of the entire iterative process.

We then searched for differences in community structure that could explain changes in leaf greenness amongst selection groups. In this aim, we first looked at microbial community structure in the selection groups between the two phases. We could not identify any effects of selection on the structure of the bacterial (*R*^2^ = 0.03, *P* = 0.82, Fig. 4, A) and fungal communities (*R*^2^ = 0.03, *P* = 0.64, Fig. 4, B) during the transitory phase (generation G01 to G05). However, when assessing the selection effect during the stabilization phase (generation G06 to G10), we detected significant effects both for bacterial (*R*^2^ = 0.08, *P* = 1.30×10^−3^, Fig. 4, C) and fungal (*R*^2^ = 0.16, *P* = 9.99×10^−5^, Fig. 4, D) communities. These results confirmed that directional selection became operational on the community structure only after stabilization was reached, and motivated the search for correlations between the evolution of all recorded plant traits and microbial community composition throughout the whole dataset. All morphological plant traits recovered from the camera-based phenotyping (convex hull perimeter, leaf area, maximum width and height, projected leaf area, density, and the greenness index; Fig. S4-S5) were considered as a “plant multivariate dataset”. Using two separate sparse partial least squares discriminant analysis, we estimated the level of congruence between the plant multivariate dataset and either the bacterial (Fig. S15) or fungal datasets (Fig. S16). We found a strong correlation between plant traits and microbial community structure (*R*^2^ = 0.61, *P* < 0.001 for bacteria, Fig. 4, D, and *R*^2^ = 0.63, *P* < 0.001 for fungi, Fig. 4, E), which can be assimilated to a causal relationship, since the transfer of microbial communities from one generation to the other was the unique source of non-random variation influencing plant traits during the experiment.

**Figure 4:**
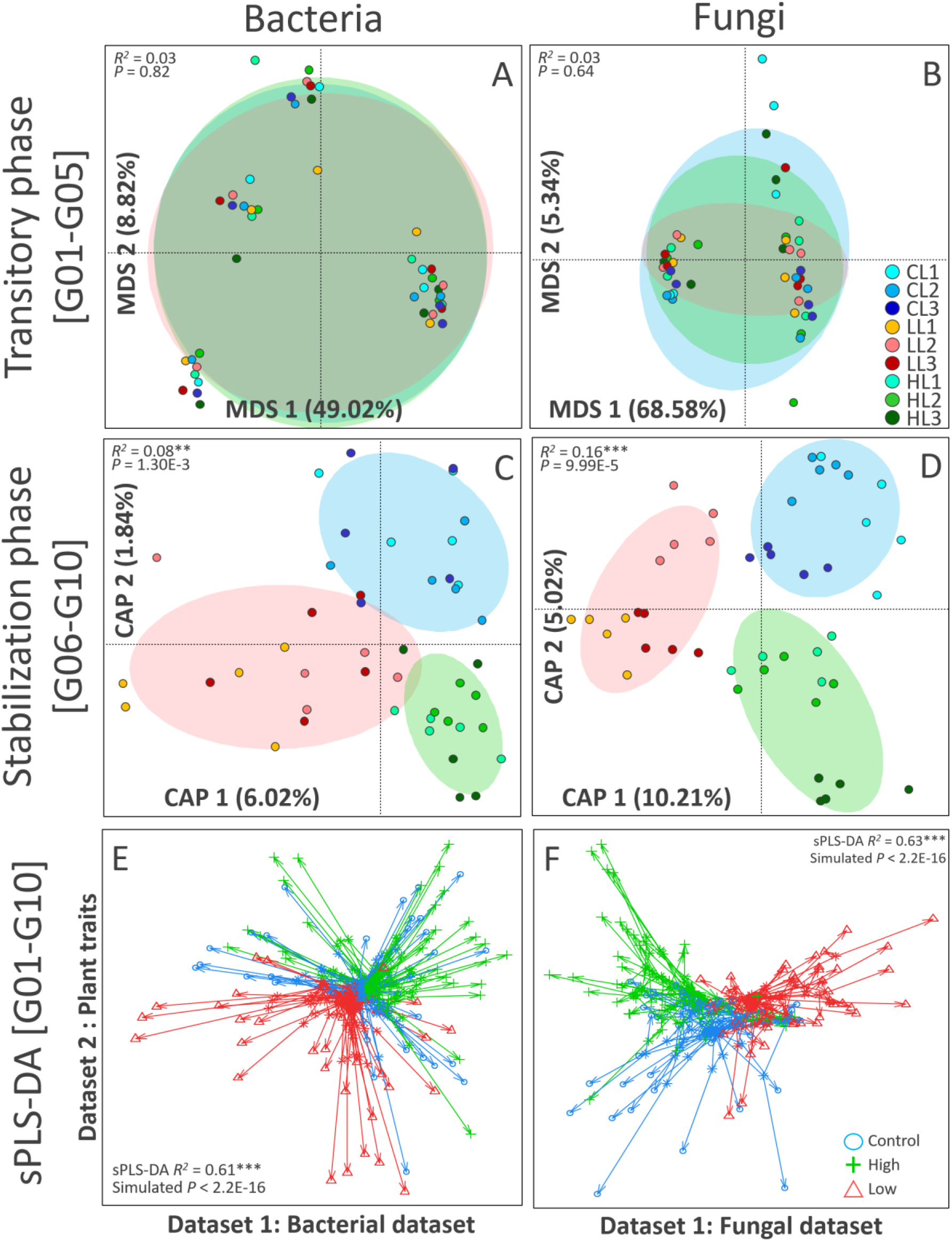
Effect of artificial selection on the selected microbiota community structure and sparse partial least square discriminant analysis (sPLS-DA). Panel A and B show the structure of the rhizosphere bacterial and fungal communities during the transitory phase [G01-G05]. Panel C and D show the structure of the rhizosphere bacterial and fungal communities during the stabilization phase [G06-G10]. The models were built using the Bray-Curtis dissimilarity index, with 10.000 group permutations (Bray-Curtis ∼ selection). The *R*^2^ values are indicating the percentage of variance explained by the model. If significant, the constrained coordinates are shown (model *P* < 0.05, CAP, Constrained Analysis of Principal coordinates). If not, the unsupervised coordinates are shown (model *P* > 0.05, MDS: Multi-Dimensional Scaling). Panel E and F are showing the results of the sPLS-DA between the plant traits dataset and the bacterial or fungal datasets, respectively. Arrow plots are showing the samples correspondence between microbial and plant data. The start of arrows indicates the location of the sample in the PCA of the dataset 1 (bacteria or fungi datasets), and the arrow tips indicate the location of the sample in the PCA of the dataset 2 (plant traits dataset). Arrow location, length and direction is corresponding to the congruence between datasets, which was tested with a randomized group simulation (N = 1,000 permutations, Fig. S15-S16).

From this analysis, we specifically excavated the microbial taxa that correlated either positively or negatively with the leaf greenness from the multivariate plant dataset, regardless of generation, selection group, or lineage. When considering only strong correlations (>|0.4|), we detected two distinct microbial sub-communities: i) one made of 325 taxa whose abundances were positively correlated with leaf greenness (hereafter called “positive taxa”, 313 bacterial and 12 fungal, Fig. S17) and ii) one made of 68 taxa whose abundances were negatively correlated with leaf greenness (hereafter called “negative taxa”, 49 bacterial and 19 fungal, Fig. S17). The taxa in the positive and negative microbial sub-communities belonged to very distinct phylogenetic groups, with a greater diversity for positive bacteria than the negative ones, while the negative fungal taxa showed slightly higher diversity than the positive ones (Fig S17). Finally, we investigated the effect of selection on the grouped relative abundance of these positive and negative microbial taxa in order to identify how they responded to directional artificial selection between the transitory and stabilization phases (Fig. S18). We noted significant changes in the relative abundance of these taxa between the two phases regardless of the selection groups, with a significant increase of positive bacterial and fungal taxa as well as a significant decrease of negative bacterial taxa and increase in negative fungal taxa (mostly explained by the low selection group, panel F) from the transitory to the stabilization phase (*P* < 0.001; Fig. S18 A-B). This general trend was well captured in the control group, which was not subjected to directional selection (blue lines and stars, control at transitory vs control at stabilization, *P* < 0.05, Fig. S18 C-F). These observations indicated that the transfer of random parental microbial communities to the offspring generations was not a neutral process, as it led to significant changes in the community structure (Fig. 4 C-D) and the abundance of taxa correlating with greenness (Fig. S18). This phenomenon, known to occur in experimental evolution experiments in the absence of selection pressure, is referred to as ‘*controlled natural selection*’^21^, and is due to the selection effect of the plant on its microbiota *via* specific recruitment mechanisms^22^. Concomitantly, directional artificial selection in the high and low selection group significantly altered the abundance of positive and negative taxa between phases. While no differences among the selection groups were observed in the transitory phase, significant effects occurred in the stabilization phase. Indeed, compared to the control group during this phase, the low selection resulted in a significant reduction of positive bacterial (red stars, from 22.44% to 14.66%, *P* < 0.01, Fig. S18, C) and fungal taxa (red star, from 8.60% to 6.35%, *P* < 0.05, Fig. S18, D), as well as a significant increase in negative bacterial (red star, from 2.05% to 2.82%, *P* < 0.001, Fig. S18, E) and fungal taxa (red star, from 1.08% to 12.93%, *P* < 0.001, Fig. S18, F). These results indicated that despite the lack of reliable effects on the selected plant property (Fig. S14), the low selection modality has resulted in a significant steering of the rhizosphere microbiota structure. On the other hand, the efficient high selection resulted in a significant increase of positive fungal taxa (green stars, from 8.60% to 13.10%, *P* < 0.001, Fig. S18, D), as well as a significant reduction of negative bacterial (green stars, from 2.05% to 1.48%, *P* < 0.001, Fig. S18, E) and fungal taxa (green stars, from 1.08% to 0.29%, *P* < 0.001, Fig. S18, E) compared to the control group during the stabilization phase. The increase of positive bacterial taxa was not significant (Fig. S18, D). Therefore, directional selection has either accelerated or slowed the controlled natural selection process instigated by the plant, by increasing or decreasing the relative abundance of phylogenetic distinct taxa correlating with the targeted property.

Directional selection of the rhizosphere microbiota is a promising strategy for modifying plant phenotypes without changing plant genotypes. Here, we provide empirical evidence that plant phenotype can be altered by exclusively transferring rhizosphere microbiota from generation to generation (Fig. 1). We observed strong oscillations in microbial community structure during the first generations, followed by the maintenance of a stable community structure (Fig. 2), with a clear breaking point at generation G05 that supported the distinction between a transitory and a stabilization phase (Fig. 3, A). Once community structure stabilized, the selected plant property became heritable between generations G06 and G10 (Fig. 3, B), concomitantly to the appearance of distinct community structures in each selection group (Fig. 4, C-D). There was a strong and significant congruence between manipulated microbial community structures and all measured plant traits, suggesting a causal effect (Fig. 4, E-F). The specific focus on microbial taxa correlating with the leaf greenness revealed significant effects in the control group between the two phases, suggesting a controlled natural selection of the plant in favor of potentially beneficial taxa in the absence of directional artificial selection (Fig. S18). Compared to the control group, we verified that the selection pressure has indeed altered the abundance of two phylogenetically distinct microbial sub-communities correlating with the property of interest. We concluded that in artificial selection of microbial communities, the heritability of the selected property depends on the stability of microbial community structure^19,20^. We believe that understanding the conditions leading to microbiota stability is an essential cornerstone for the development of efficient microbiota selection programs, that deserves increased attention in future research in this field.

## Acknowledgments

We would like to thank the members of the 4PMI platform for their expertise and help during plant phenotyping (For Plant and Microbe Interaction, INRAE Centre Dijon, France, https://www6.dijon.inra.fr/umragroecologie/Plateformes/Serres-PPHD). Respectively, we thank Damien Gironde, Frédéric Saignole, Noureddine El-Mjiyad and Karine Palavioux for helping during plant growth monitoring; Franck Zenk and Julien Martinet for image capture; and Mickael Lamboeuf for image processing; Sébastien Anselme, Richard Sibout and Thomas Girin from the *Brachypodium* resources center at the Institut Jean Pierre-Bourgin, (INRAE Centre Versailles, France) for seeds provision; Beatriz Decencière, Amandine Hansart and Florent Massol of the CEREEP - Ecotron IDF/UMS CNRS/ENS 3194 for soil provision. We would like to thank Tiffany Raynaud, Luiz Domeignoz Horta, Florian Bizouard, Eric Pimet and Chantal Ducourtieux for their technical help during soil sampling, the artificial selection and the post-selection experiments. We would like to thank Annick Matejicek for her help with the CHN measures. We also thank Jessica Pearce-Duvet for the English editing of our manuscript.

## Author contributions

Conceptualization, funding acquisition and project administration (MB). Supervision and methodology (MB and SJ). Investigation (SJ, SW, VM, DB). Resources (4PMI: CS, molecular biology: LP, sequencing: SJS). Formal analysis (SJ and AS). Validation (AS, LP and MB). Writing - original draft (SJ and MB). Writing – review & editing (AS, SW, VM, DB, SJS, CS, LP).

## Data and materials availability

Sequencing data generated in this study has been deposited in the Sequence Read Archive public repository (SRA: https://www.ncbi.nlm.nih.gov/sra/docs/) with an embargo of six months or until publication of the manuscript (Accession number for the bacterial and fungal datasets: SUB7720753 and SUB7738355). Images of plants are stored at the 4PMI platform server (350973 files worth 463gB of data) and can be made available upon request addressed to the corresponding author. The Matlab RGB routine script used at the 4PMI platform to estimate leaf colors can be made available upon request to the corresponding author. Plant traits data is available as supporting data. Statistical analysis was performed with the publically available software Rgui, with function packages and documentation that are publically available and referenced. The Rgui scripts can be made available upon request to Samuel Jacquiod.

## List of Supplementary Materials

- Materials and Methods (see Supplementary Materials file)
- Figs. S1 to S19
- Table S1 to S2
- References that are only cited in Supplementary Materials: [22-44]

