## Supplementary material for "Artificial selection of stable rhizosphere microbiota leads to heritable plant phenotype changes": materials and methods

### Title

**This section includes:**

- Materials and Methods
- Figs. S1 to S19
- Tables S1 to S2

**Materials and Methods**

### i. Soil

An oligotrophic sand soil was selected to facilitate rhizosphere sampling and reinforce the plant-microbe nutrition interaction (cambisoil with moor; organic carbon: 14.7 g $kg^{-1}$; total nitrogen: 1.19 g $kg^{-1}$; pH: 5.22; sand: 74.1%, loam: 19.0%, clay: 6.9%, origin: 48°16'58.9"N 2°40'19.0"E, CEREEP Ecotron, Saint-Pierre-Lès-Nemours, France[23]). Approximately two tons of fresh bulk soil freed from plant material were sampled and immediately transported to the 4PMI platform (Plant Phenotyping Platform for Plant and Microorganisms Interactions, Agroécologie, INRAE Centre Dijon, France). The soil was dried for two weeks with daily mixing at ambient temperature (T° = 20±5°C) and sieved at 2mm, yielding ~850 kg which were stored in a sealed container box. Before each generation batch, 65 kg of soil was autoclaved once (120°C, 40 min) and allowed to rest for 48 h. A total of 180 pots were prepared at each generation, containing 350 g of soil each, and equipped with individual cups for sub-irrigation. 80 ml of osmosed water was added per pot to reach 90% of the water holding capacity (206 mg water per gram of dry soil). Pots were set at room temperature for 24 h before seedling transplantation. After seedling transplantation, pots were randomly distributed on 16 trays and placed in the climatic chamber.

### ii. *Brachypodium distachyon* growth condition

The model plant for cereals, *Brachypodium distachyon* Bd21 (wild type), was selected for its convenient small size and available large documented knowledge concerning its morphology and physiology[24-26]. For all generations, we used the same seed batch provided by the Institut Jean-Pierre Bourgin (Versailles, France). For each generation, ~300 seeds were germinated in boxes containing blotting paper saturated with osmosed water, placed in a dark cold chamber (4°C, 24 h), moved to a germinator (Fitoclima 600 PL/PLH, Aralab) in the dark (18°C, 48 h), and then exposed to full light (21°C, 72 h). Homogeneous seedlings were randomly assigned to the 180 pots and moved to the climatic chamber for four weeks in random position (21°C, 10 h light per day, 70% air humidity, 200 $\mu mol.s^{-1}.m^{2}$ light intensity, Walk-in EH, Aralab light intensity). Pots were watered every second day with fixed doses and once a week, all pots were weighted individually to be reset at 90% of the water holding capacity. The position of pots within trays, and trays themselves were randomly redistributed in the chamber once a week.

### iii. Initial microbial pool and inoculation of the first generation

Once in pot, the soil from the first generation was inoculated with an initial microbial pool obtained from soil adhering to the roots (at 70% humidity) of three *Brachypodium distachyon* plants grown in the same soil, but not autoclaved, for five weeks (Fig. S2). Rhizosphere soil was mixed with 600 ml osmosed water, and stirred for 30 min with a magnet-bar mixing to obtain a liquid slurry that was used as our initial inoculation pool. Thereafter, the 180 transplanted seedlings of the first generation were inoculated with this liquid slurry. Part of the slurry was centrifuged (8000 rpm, 10 min) and split in half for i) glycerol storage at -80°C (25% final) and ii) DNA extraction.

### iv. Plant phenotyping and selection based on image analysis

Plants and their associated rhizosphere microbiota were selected based on a non-destructive procedure to preserve live plants and associated microbial communities (Fig. S2). For this purpose, we defined a leaf greenness index, a proxy health and nutrition status[8], based on dark green coloration of *Brachypodium distachyon* shoot images acquired at the 4PMI platform (https://www6.dijon.inrae.fr/plateforme4pmi_eng/Technical-description/High-Throughput-phenotyping, Fig. S1). After four weeks of growth in the climatic chamber, plants were taken out and sent to the automatic plant phenotyping device available at the 4PMI platform for image acquisition, consisting of twelve side-views at different angles in a light-controlled cabin for each plant (Fig. S4). Images were analyzed *via* the routine pipeline available at the 4PMI platform[27,28], enabling the estimation of several morphological parameters being: picture shooting time, maximum height over-pot, maximum width, convex-hull perimeter, convex-hull area, projected leave surface area (object), density and colored surfaces (Fig. S4, B, C and D). We specifically quantified the surface of leaves with dark green pixels and established a leaf greenness index as a ratio between the desired colored surface (dark green) multiplied by the total plant surface (Object surface) and divided by the remaining surface (Object surface – dark or light green surface, Fig. S4, D). The formula was set as follow:

$$\text{Leaf greenness index} = (\text{Green surface} * \text{Object surface}) / (\text{Object surface} - \text{Green surface})$$

Multiplying by the whole "Object surface" was added to avoid selecting small plants with high greenness ratio due to nutrient accumulation in lesser biomass. Therefore, at the end of the growth phase of each generation, shoot images of the 180 experimental pots were acquired (2160 images per generation) and analyzed to define a leaf greenness index for each plant that was used as our artificial selection criterion.

#### v. Statistical analysis of the leaf greenness index and other plant traits

Statistical analyses of the camera-based plant traits were performed using the Rgui software[28]. For the leaf greenness, normality was assumed after visual inspection of the residual variance normality and validated with the d'Agostino test to rule out skewness in the residuals ($P > 0.01$, package '*moments*'[30]). If normality of the data could not be confirmed, non-parametric tests were used. To be in line with our hypothesis that directional selection will result in a significant increase or decrease of plant traits compared to the random selection, statistical analysis of the high and low selection groups were always done against the random control group via one-sided version of the statistic tests (Welch t-test if normal; Wilcoxon Rank Sum test if not, alternative = "greater" for the high selection group, and "less" for the low selection group). Out of the 1800 *Brachypodium distachyon* grown during the selection on the sand soil (180 plants x 10 generations), 21 plants were not included due to death or too low growth for proper image analysis (leaf greenness index < 10).To account for the predominant generation effect, we used the control selection group (random selection) to standardize the greenness values of plants in the high and low selection groups using the z-score normalization. Briefly, we respectively subtracted and divided the individual plant leaf greenness values within a given generation by the average and standard deviation of all individuals of the three control lineages of that same generation.

#### vi. Experimental protocol of the selection experiment

For the generations 1 to 10, and out of the 20 plants available within each lineage, we selected either the three plants with the highest leaf greenness for the high selection modality, or the plants with the lowest leaf greenness for the low selection modality (Fig. S2). For the control lineages, we randomly drew numbers without replacement between 1 and 20 to select the plants.

We carefully dismantled the root systems of selected pots to recover the adhering soil. To rule out the eventual rhizosphere biomass bias that could arise from the directional selection of large or small plants, we systematically normalized the sampling step according to the adhering soil weight, starting with the three smallest rhizospheres obtained in each of the low selection lineages to calibrate the target weight of material recovered next in the high and control selection, which had bigger root systems. This procedure resulted in selection of pooled rhizosphere microbiota with stable molecular abundances throughout the entire selection procedure, as confirmed *a posteriori* by quantitative PCR of the 16S rRNA gene and ITS2 markers, showing only marginal levels of fluctuations below 1 order of magnitude (Fig. S9, see section ix.). For the dismantling, soil humidity was set to 70% WHC to ensure a sufficient amount of rhizosphere soil adhering to the roots after gentle shaking. For each lineage, the three selected root systems and tightly adhering rhizosphere soil were pooled and mixed up with 200 ml of osmosed water, and stirred 30 min with a magnet-bar mixing (Fig. S3). Pooling of several rhizosphere was applied to ensure enough microbial diversity and variability to apply artificial section in satisfactory conditions[7]. 100 ml of the recovered slurry was centrifuged in two 50 ml falcon tubes (8000 rpm, 10 min). One falcon tube was used to save the microbial communities in glycerol at 30% final concentration (-80°C), and the other half was saved for DNA extraction (-20°C). The remaining 100 ml of slurry were used to inoculate the next seedling generation. One-week old seedlings coming from the germination boxes were directly transplanted and inoculated into the pots (Fig. S3). All pots were set in a climatic chamber according to previously described growth conditions (ii.).

vii. <u>Sampling of the offspring plant rhizosphere at the las generation</u>

At the end of the selection experiment (generation 10), six pots with homogeneous plant development (excluding the three best or worse plants in terms of greenness) were selected within each lineage (6 x 9 = 54), and their individual rhizosphere soils were recovered to assess the effect of artificial selection on individual rhizosphere microbial communities. Each individual root system and tightly adhering soil was vortex-washed in a 50ml falcon tube (1min, 5ml osmosed water) and the recovered soil slurry was frozen at -20°C.

viii. <u>Microbial community analysis</u>

Rhizosphere soil samples and inoculation pools were analyzed for total bacterial and fungal diversity and composition by sequencing respectively the 16S rRNA gene (small subunit of the prokaryotic ribosomal operon) and ITS2 region (Internal Transcribed Spacer 2 in the ribosomal operon of fungi) via Illumina Miseq 2 x 250 bp paired-end analysis. In total, 145 samples were extracted, including: i) 91 inoculation pools generated during the selection experiment (initial diversity pool + 9 lineages x 10 generations) and ii) 54 individual rhizospheres sampled in generation 10 (6 rhizosphere x 9 lineages). First, DNA was extracted from 250 mg of rhizospheric soil slurry using the DNeasy PowerSoil-htp 96 well DNA isolation kit (Qiagen, France). 16S rRNA gene and ITS amplicons were generated for all extracts in two steps. In the first step, the V3-V4 hypervariable region of the bacterial 16S rRNA gene was amplified by polymerase chain reaction (PCR) using the fusion primers U341F (5’-CCTACGGGRSGCAGCAG-3’)[31] and 805R (5’-GACTACCAGGGTATCTAAT-3’)[32], with overhang adapters (forward: TCGTCGGCAGCGTCAGATGTGTATAAGAGACAG, adapter: GTCTCGTGGGCTCGGAGATGTGTATAAGAGACAG) to allow the subsequent addition of multiplexing index-sequences. Note that U341F and 805R also target Archaea sequences, but since those were extremely minor in our datasets (~0.01%), we considered amplicon generated with those primers as belonging to bacteria. Fungal ITS2 was amplified using the primers gITS7F (5'- GTGARTCATCGARTCTTTG-3')[33] and ITS4R (5'-TCCTCCGCTTATTGATATGC-3')[34]. Thermal cycling conditions of the first step PCR were as follows: 98°C for 3 min followed by 98°C for 30 sec, 55°C for 30 sec and 72°C for 30 sec (25 and 30 cycles for 16S rRNA and ITS genes, respectively) and a final extension for 10 min at 72°C. Duplicate first step PCR products were pooled and then used as template for the second step PCR. This second PCR amplification added multiplexing index-sequences to the overhang adapters using a unique multiplex primer pair combination for each sample. Thermal cycling conditions were as follows: 8°C for 3 min followed by 98°C for 30 sec, 55°C for 30 sec and 72°C for 30 sec (8 and 10 cycles for 16S rRNA and ITS genes, respectively) and a final extension for 10 min at 72°C. Duplicate second step PCR products were pooled then visualized in 2% agarose gel to verify amplification and size of amplicons. Amplicon products were purified using HighPrep™ PCR Clean Up System (AC-60500, MagBio Genomics Inc., USA) paramagnetic beads using a 0.65:1 (beads:PCR reaction) volumetric ratio to remove DNA fragments below 100 bp in size and primers. Samples were normalized using SequalPrep Normalization Plate (96) Kit (Invitrogen, Maryland, MD, USA) and pooled using a 5 µl volume for each sample. The pooled samples library was concentrated using DNA Clean and Concentrator™-5 kit (Zymo Research, Irvine, CA, USA). The pooled library concentration was determined using the Quant-iT™ High-Sensitivity DNA Assay Kit (Life Technologies). Before library denaturation and sequencing, the final pool concentration was adjusted to 4 nM. Amplicon sequencing was performed on an Illumina MiSeq platform using Reagent Kit v2 [2 x 300 cycles] (Illumina Inc., CA, US). Demultiplexing and trimming of Illumina adaptors and barcodes was done with Illumina MiSeq Reporter software (version 2.5.1.3).

16S rRNA gene and ITS amplicon sequences were analyzed in-house with a Python notebook (available upon request). For consistency reason in the way we have analyzed our 16S rRNA gene and ITS2 amplicon, we have decided to use a well-established OTU pipeline instead of Amplicon Sequence Variants (ASV) since the latter is currently not available for fungal ITS. Briefly, 16S rDNA and ITS sequences were assembled using PEAR[35] with default settings. Further quality checks were conducted using the QIIME pipeline[36] and short sequences were removed (< 400 bp for 16S and < 300 bp for ITS). Reference based and *de novo* chimera detection, as well as clustering into operational taxonomic units (OTUs, hereafter called taxa) were performed using VSEARCH[37] and the adequate reference databases (*Silva* representative set of sequences for 16S rRNA and UNITE's ITS2 reference dynamic dataset for ITS). The identity thresholds were set at 94 % for 16S rRNA and 97 % for ITS amplicon based on a routine mock community used as an internal calibration. Taxonomy was assigned using UCLUST[38] and the latest release of *Silva* database (v138.1)[39]. For ITS, the taxonomy assignment was performed using BLAST algorithm and the UNITE reference database (v.7-08/2016)[40]. Raw sequences have been deposited to the SRA public repository (Sequence Read Archive, 16SrRNA), with an embargo period of 6 months or until publication

ix. Real-time quantitative PCR (qPCR)

The molecular abundances of total bacterial and fungal microbial communities in selection pools of all lineages for all generations were estimated by real-time quantitative PCR (qPCR). Total bacterial and fungal communities were quantified using 16S rDNA and ITS primers[34,41]. qPCR reactions were carried out in a ViiA7 (Life Technologies, USA) in a 15 µl reaction volume containing 7.5 µL of Takyon Master Mix (Eurogentec, France), 1 µM of each primer, 250 ng of T4 gene 32 (QBiogene, France) and 1 ng of DNA. Two independent runs were performed for each real time PCR assay. Standard curves were obtained using serial dilutions of linearized plasmids containing appropriated cloned targeted genes from bacterial strains or environmental clones. PCR efficiency for the different assays ranged from 88 to 113 %. No template controls gave null or negligible values. Inhibition in qPCR assay was tested by mixing soil DNA extracts with either control plasmid DNA (pGEM-T Easy Vector, Promega, France) or water. No inhibition was detected in any case.

x. Analysis of microbial communities of the pools during the course of selection

A summary of the different samples sequenced is presented in Table S2. To account for uneven sequencing depth amongst samples, rarefaction curves were calculated with the package '*vegan*'[42] to assess sequencing depth before random resampling to normalize samples in each dataset (Fig. S19). The inoculation pools from the selection experiment (n = 91 samples) were rarefied at 45,000 and 17,000 sequences per sample for bacteria and fungi, respectively. The last generation (G10) datasets (n = 54 samples) were rarefied at 17,000 counts together with the selection datasets. A level of 10,000 is considered to be optimal for alpha-diversity estimation[43].

Alpha-diversity analysis was performed to follow the status of rhizosphere microbiota during selection using the following indices: observed taxa richness (S), Simpson index (1-D, D = Dominance), Shannon index (H) and Equitability (E). Alpha-diversity indices were exported for bacteria (Fig. S7) and fungi (Fig. S8). Bacterial and fungal molecular abundances were verified by qPCR, showing relatively stable levels throughout the entire selection experiment across generations (Fig. S9). Log10-models were adjusted, and their significance validated by verifying the linearity between fitted and observed values ('*cor.test*', $P < 0.001$). For the beta-diversity analysis, the Bray-Curtis dissimilarity index was used to generate distance matrices for both bacterial and fungal datasets. For consistency reasons, the Bray-Curtis dissimilarity index was preferred over other metrics (e.g. UniFrac) as ITS sequences cannot be aligned to obtain a phylogenetic distance. A distance-based redundancy analysis (db-RDA) was used with the following model: distance ~ generation*selection/lineage ('*capscale*' function, package '*vegan*', 10,000 permutations, Fig. 2, A-B). The refined analysis of the selection effect was done for each phase with the model: distance ~ selection ('*capscale*' function, package '*vegan*', 10,000 permutations, Fig. 4, A-D)

xi. Analysis of the offspring rhizospheres obtained at the end of the selection

For the last generation dataset (G10), the purpose was to estimate the evolution of rhizosphere microbiota dissimilarity overtime by comparing six individual rhizospheres of each lineage from the last generation (G10) with all the consecutive inoculation pools of that same lineage (Fig. S10, A). The selection pools were also included and rarefied to the same level based on the lowest sample (bacteria = 17,000; fungi = 17,000). The Bray-Curtis dissimilarity index was used to estimate, within each lineage, the evolution of microbiota beta-diversity during the selection by pairwise comparisons (Fig. S10, A). Beta-diversity evolution of bacterial and fungal communities was modelled using an unsupervised segmented analysis to determine the presence of an eventual breaking point during selection in each lineage independently (Fig. S11-S12), and on all the lineages at once (Fig. 3, A) (package '*segmented*')[44].

xii. Sparse partial least square discriminant analysis (sPLS-DA)

We aimed to identify microbial taxa associated with alteration in plant traits (greenness, height, width, density, perimeter, area…), by using the sparse partial least square discriminant analysis (sPLS-DA) implemented in the '*mixOmics*' package (function '*block.splsda'*)[45]. This method is specifically designed for integration of several multivariate datasets generated from the same samples in order to identify correlated variables explaining a categorical outcome (http://mixomics.org/mixdiablo/). Two independent sPLS-DA models were generated, one for bacteria and the other for fungi. The number of components to integrate in the analysis was calculated to minimize the error rate *via* a performance diagnostic (N = 8 for the bacterial model, N = 5 for the fungal model, function '*perf.diablo*')[45]. The congruence between the plant traits and microbiota community structure was assessed based on diagnostic plots and a random simulation with 1.000 group permutations to test the significance of the selection procedure on the first two main components of each model (Fig. S15-S16). We specifically extracted from the sPLS-DA models the taxa whose abundance was correlating the leaf greenness index by filtering weak associations to focus on strong ones (|correlation strength| > 0.4). Two categories of taxa were made: i) the microbial taxa correlating positively with greenness, and ii) the microbial taxa correlating negatively with greenness. The phylogenetic composition of the retained taxa is presented in Fig. S17, and their grouped abundance was displayed for each category and each microbial group for the transitory and stabilization phases (Fig. S18).

**Fig. S1**

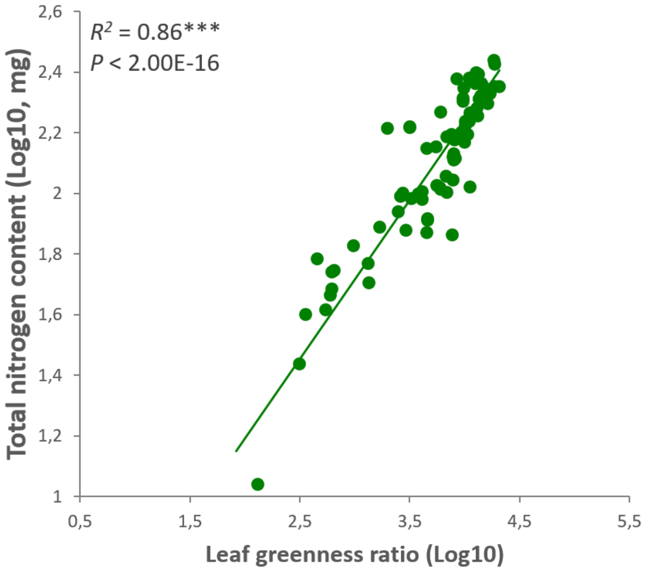


**Fig. S1: Correlation between the leaf greenness ratio index and total plant nitrogen content.** Linear relationship between the leaf greenness index obtained from camera-based phenotyping (x) and actual measured nitrogen content in *Brachypodium distachyon* shoots (y). The linear estimation was carried on dark green surfaces (y = 0.5254x + 0.1435, adjusted $R^2$ = 0.86, $P$ < 2.0E-16). Data were obtained from a batch of *Brachypodium distachyon* grown in a climatic chamber at the 4PMI greenhouse facility (INRAE BFC Centre Dijon, France) with varying $KNO_3$ fertilization to span variable nitrogen content in shoots for accurate estimation of the relationship with the leaf greenness (N = 80 replicates).

**Fig.S2.**

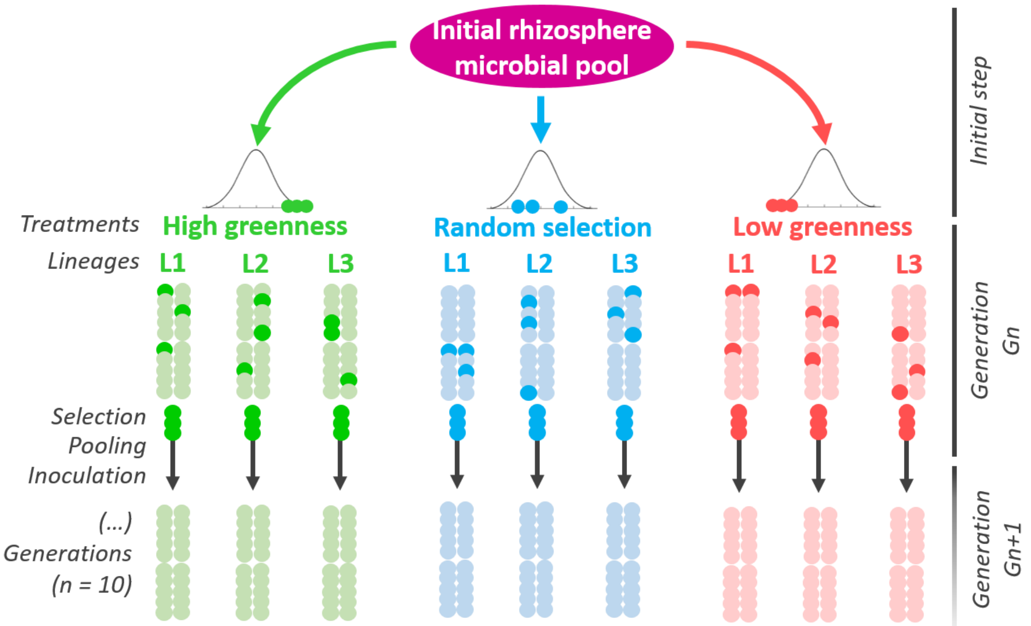


**Fig. S2: Experimental design of the artificial selection experiment**. In the first generation, all pots were inoculated with an initial microbial pool obtained from a liquid extraction of three rhizospheres of *Brachypodium distachyon*. Three distinct artificial selection groups were applied: "High selection", "Random selection" or "Low selection". In the "High" and "Low" modalities, plants with the highest or lowest leaf greenness scores (directional selection) were respectively selected. For the "Random" modality (control group), plants were picked using random drawings and served as our selection control to test if the significance of the experimenter's choice. To test the reproducibility of artificial selection, we set up three independent lineages within each selection modality (L1, L2, L3). Each lineage consisted in a population of 20 plant individuals (biological replicates) within which artificial selection was applied (e.g. in each high lineage, we selected only for the highest leaf greenness scores). Three plants were selected within each lineage to prepare the inoculum for the next generation to ensure enough genetic mixing and diversity. Once sampled, the rhizosphere microbial communities were pooled and inoculated to a new batch, with 20 plants in each lineage. This procedure was iteratively repeated for 10 consecutive "generations".

**Fig. S3.**

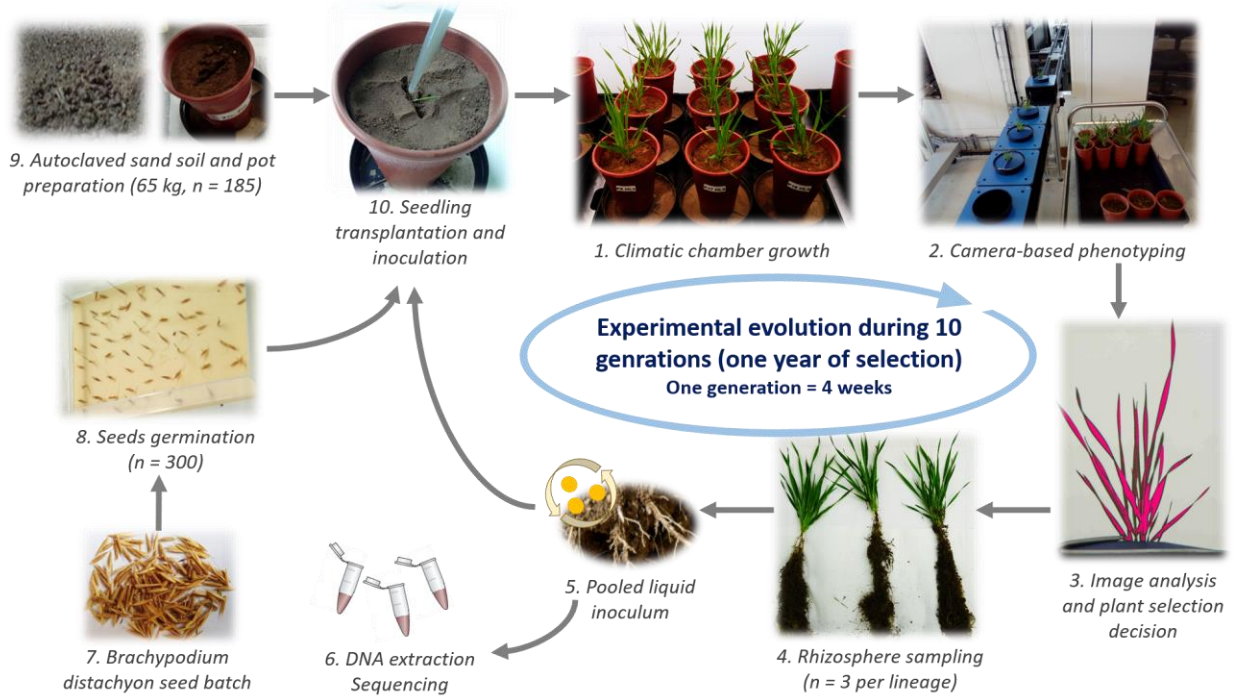


**Fig. S3: Detailed workflow of a typical generation.** Each generation consisted in a growth phase of four weeks in a climatic chamber (1), followed by a non-destructive camera-based plant phenotyping (2) to determine the level of leaf greenness (3) and select the adequate individuals (4). The rhizospheres of chosen plants were sampled and pooled to obtain a liquid inoculum (5). Part of the inoculum was saved for DNA extraction (6). Meanwhile, a new generation of seedling were prepared from the same seed batch (7-8). One week old seedlings were transplanted to new pots (9), inoculated with the liquid slurry (10), and immediately placed into the climatic chamber for another cycle of growth (1).

**Fig. S4.**

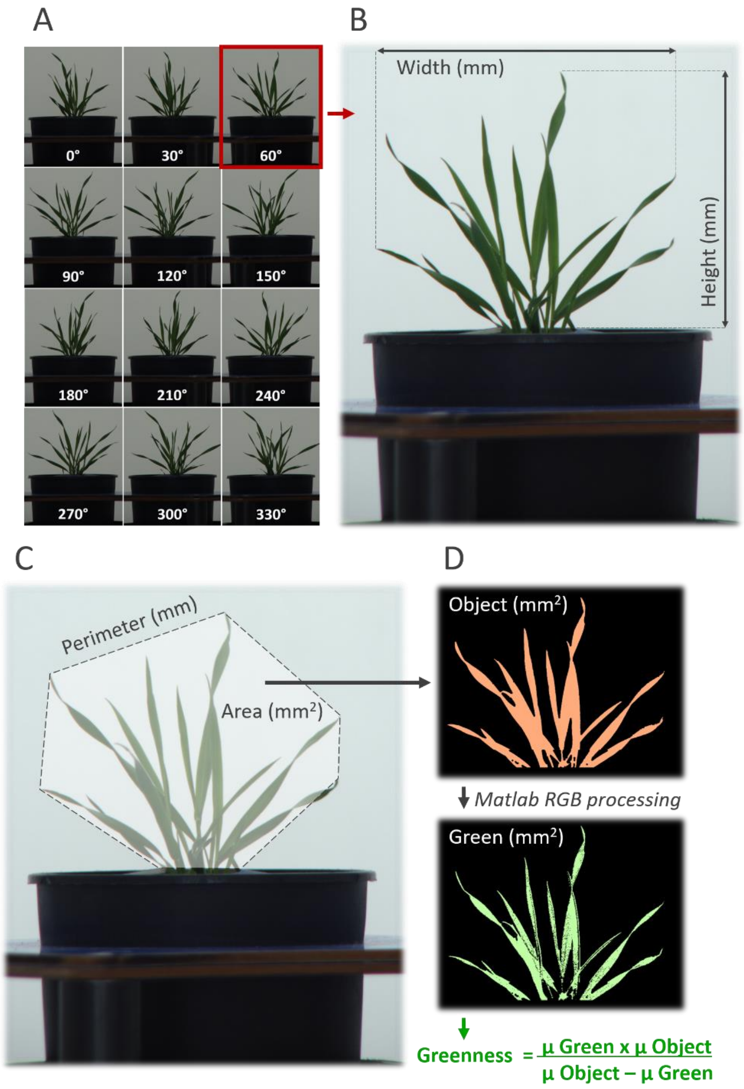


**Fig. S4: Camera-based plant phenotyping**. This figure describes the picture-based plant phenotyping procedure applied at the 4PMI platform (INRAE Centre Dijon, France). For each plant, 12 side-pictures were taken at different angles (30° rotations, A). For each picture, we estimated the maximum height over pot and width (B, example on picture at 60°), as well as the perimeter and area of the convex hull (C). For each picture, the projected leaf surface area was extracted from the convex hull area (Object, D), and we applied pixel binning using RGB hue thresholds in Matlab. Pixel values were transformed into millimeters using the picture of a standard target panel used for each generation. The leaf greenness index used for selection was calculated from the averaged values of projected leaf surface area (Object) and green surface.

**Fig. S5.**

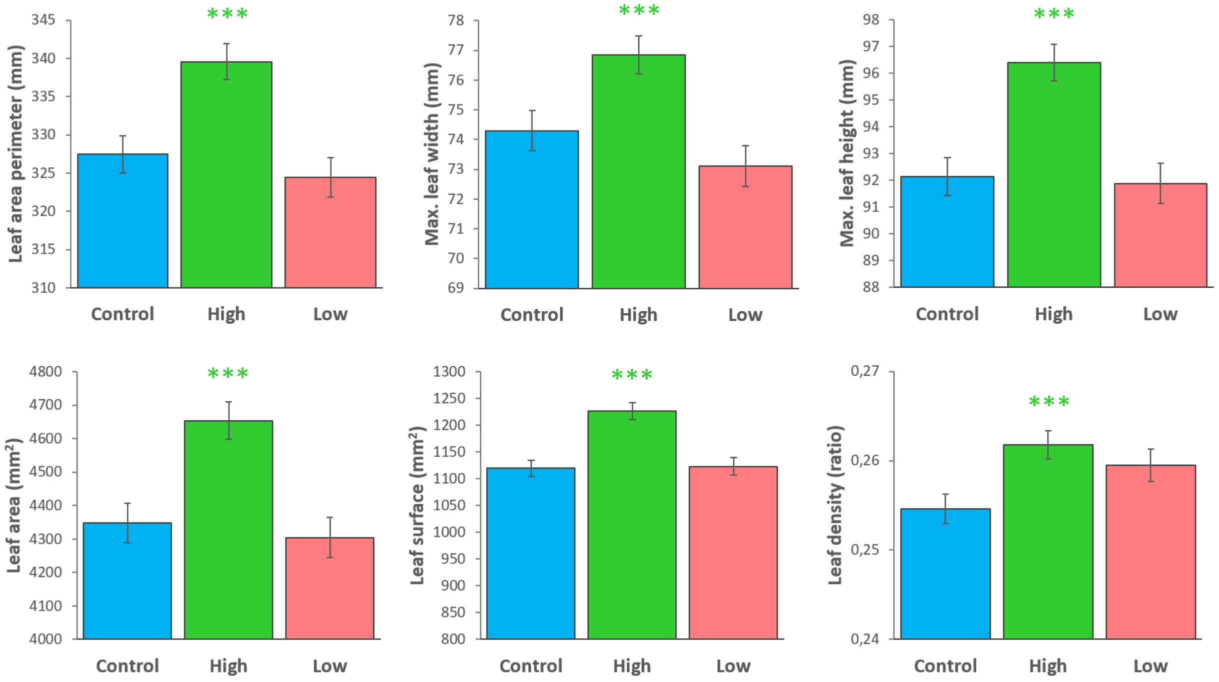


**Fig. S5:** Other morphological plant traits recorded from the camera-based phenotyping platform. Statistical comparison were done against the control group with a one-sided Wilcoxon rank sum test. *P*-value significance: « *** » $P < 0.001$; « ** » $P < 0.01$; « * » $P < 0.05$; « . » $P < 0.1$. N = 591-596 replicates for each selection group. Error-bar are representing the standard error of the mean.

**Fig. S6.**

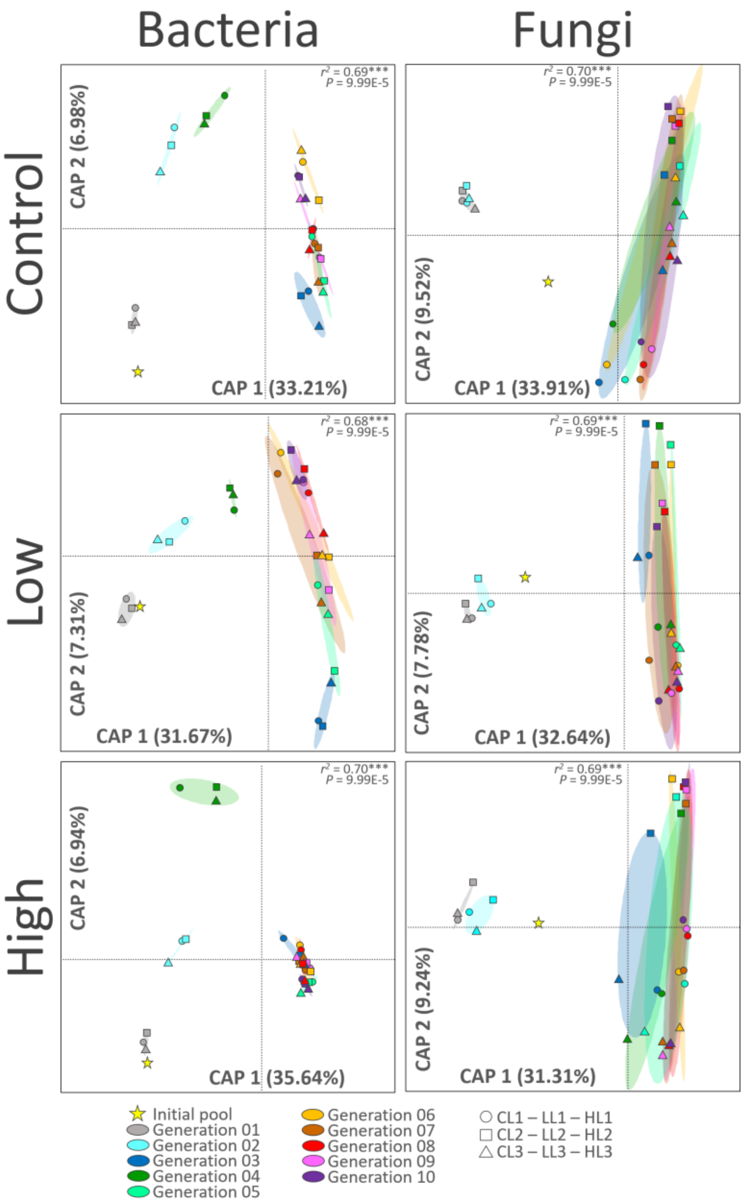


**Fig. S6:** Distance-based redundancy analysis of the selected microbiota rhizosphere for each selection group. The models were built using the Bray-Curtis dissimilarity index, with 10.000 group permutations (Bray-Curtis ~ generation/lineage). The $R^2$ values are indicating the percentage of variance explained by the model. If significant, the constrained coordinates are shown (model $P < 0.05$, CAP, Constrained Analysis of Principal coordinates). If not, the unsupervised coordinates are shown (model $P > 0.05$, MDS: Multi-Dimensional Scaling).

**Fig. S7.**

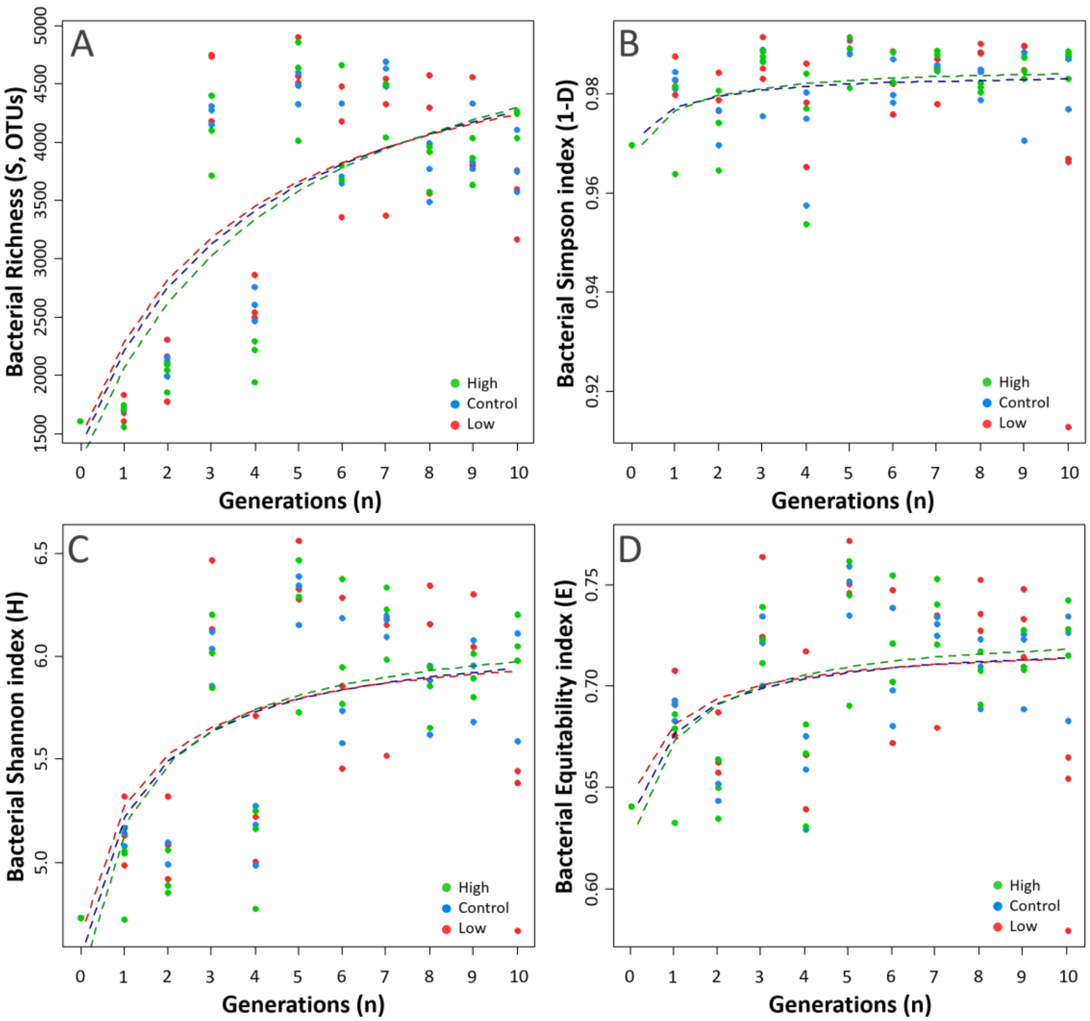


**Fig. S7: Diversity of bacterial communities across generations**. Figure shows respectively the observed bacterial taxa richness index “S” (A), the Simpson index “1-D” (B), the Shannon index “H” (C) and the Equitability index “E” (D). Alpha-diversity indices were calculated on the lineage inoculation pools used for each generation. The significance of Log-models was verified by testing the accuracy between predicted and observed values (FDR-corrected $P < 0.05$). Dotted-lines were plotted only if significance was achieved.

**Fig. S8.**

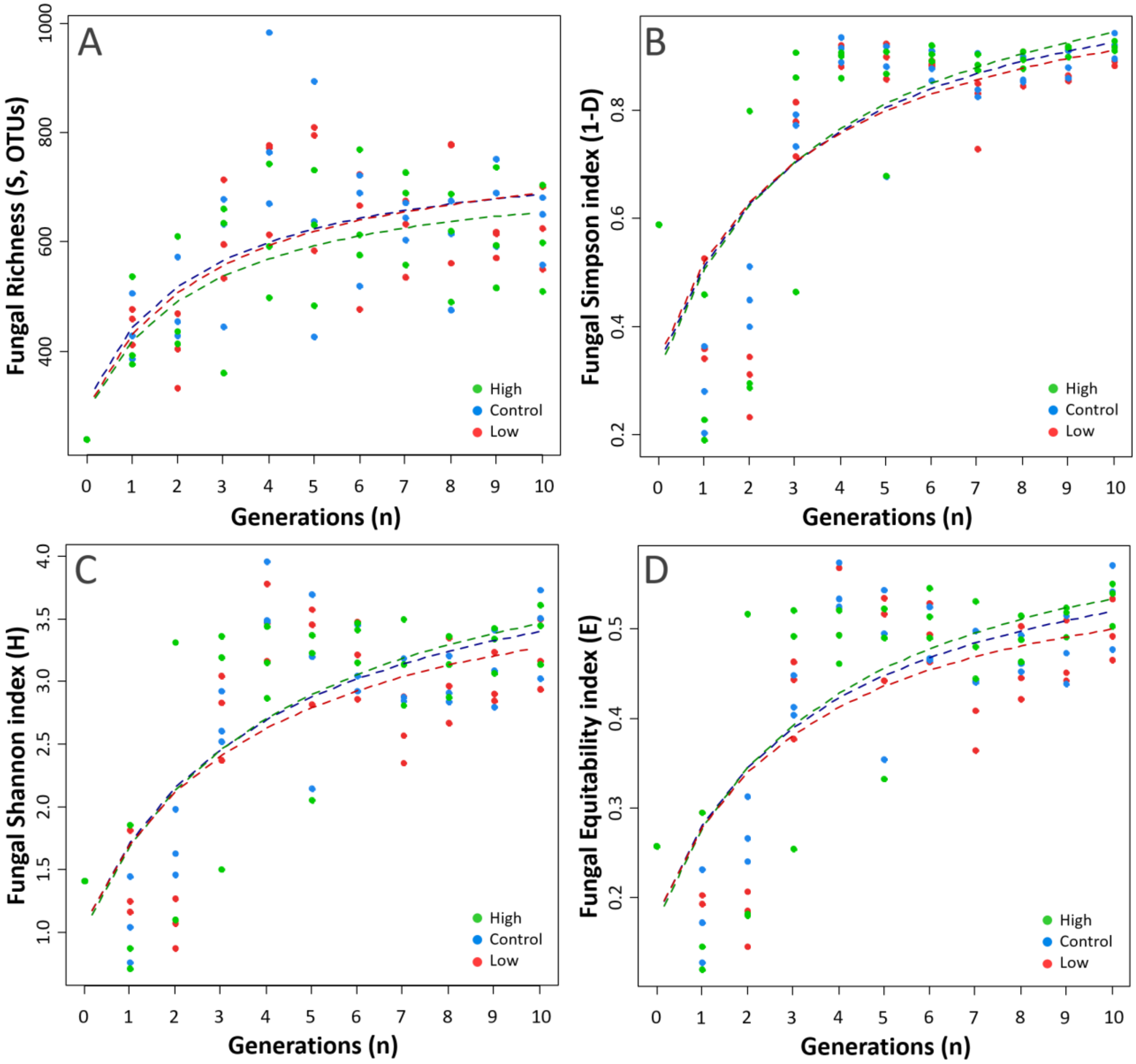


**Fig. S8: Diversity of fungal communities across generations**. Figure shows respectively the observed bacterial taxa richness index "S" (A), the Simpson index "1-D" (B), the Shannon index "H" (C) and the Equitability index "E" (D). Alpha-diversity indices were calculated on the lineage inoculation pools used for each generation. The significance of Log-models was verified by testing the correlation between expected and observed values (FDR-corrected $P < 0.05$). Dotted-lines were plotted only if significance was achieved.

**Fig. S9.**

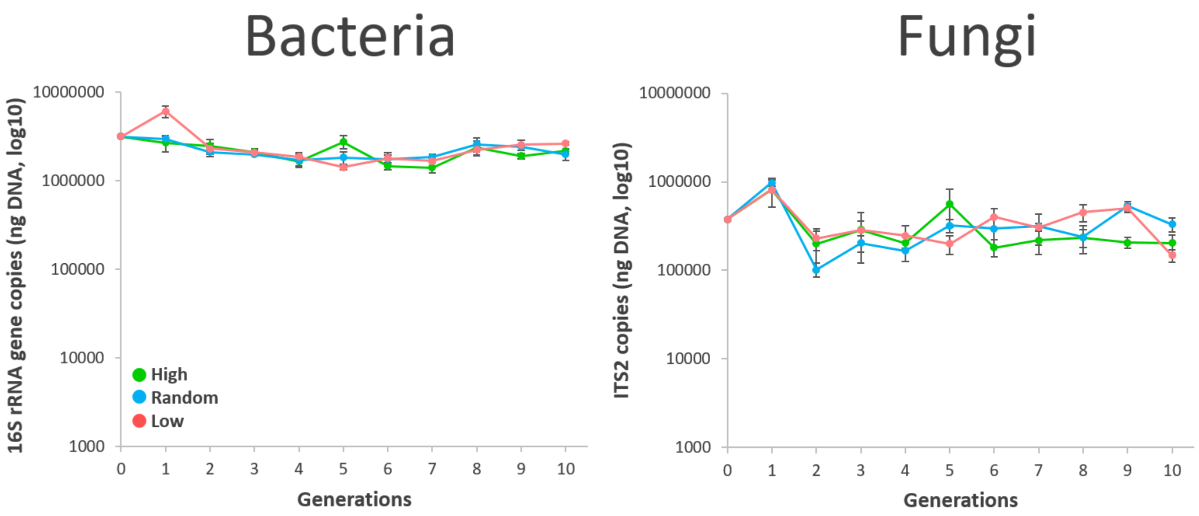


**Fig. S9:** Molecular quantification of selected bacterial (16S rRNA gene fragments) and fungal (ITS2 fragments) communities across all generations. Each point is the average between the three lineages for each selection group. Error-bars are the standard error of the mean.

**Fig. S10.**

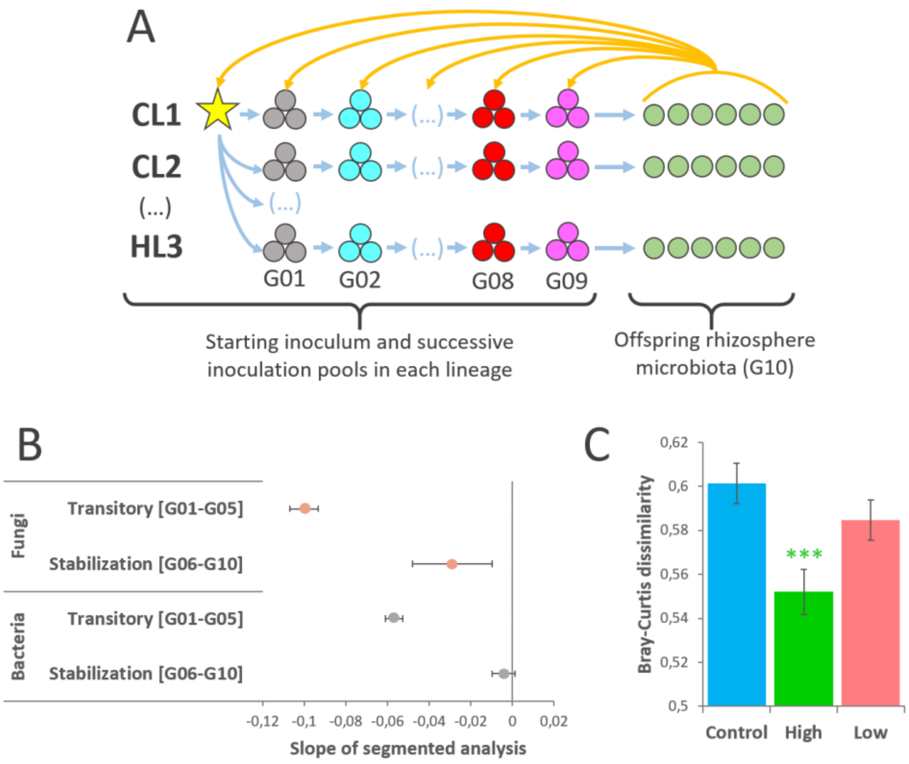


**Fig. S10:** Workflow and main results of the beta diversity analysis applied at the lineage level (Bray-Curtis dissimilarity). Panel A shows the method used to study the evolution of the beta diversity within each lineage, by respectively comparing six individual offspring rhizospheres obtained at the end of the selection procedure (G10) to each of their parent pools in descending order until the initial starting inoculum used to seed the entire experiment (G09, G08, …, G01, starting inoculum, the yellow star). The results of this analysis are further presented in supporting data for the bacterial (Fig. S11) and the fungal lineages (Fig. S12). Panel B shows the results of the unsupervised segmented linear analysis applied to the overall bacterial and fungal lineage dataset presented in figure 3, A. This panel shows the 95% confidence intervals of the slopes, before and after the breaking point at generation G05, displaying a clear stabilization for bacteria (slope not significantly different from zero from [G06-G10]), will the fungal communities are still decreasing from [G06-G10] but at a much slower pace compared to [G01-G05]. Panel C shows the overall beta-diversity average obtained in each selection group, showing that lineages from the high selection group had significantly lower dissimilarity scores compared to the control group and the low selection (two-sided Wilcoxon Rank Sum test, $P < 0.001$, N = 396 for each group, Error-bar are representing the standard error of the mean). *P*-value significance: « *** » $P < 0.001$; « ** » $P < 0.01$; « * » $P < 0.05$; « . » $P < 0.1$.

**Fig. S11.**

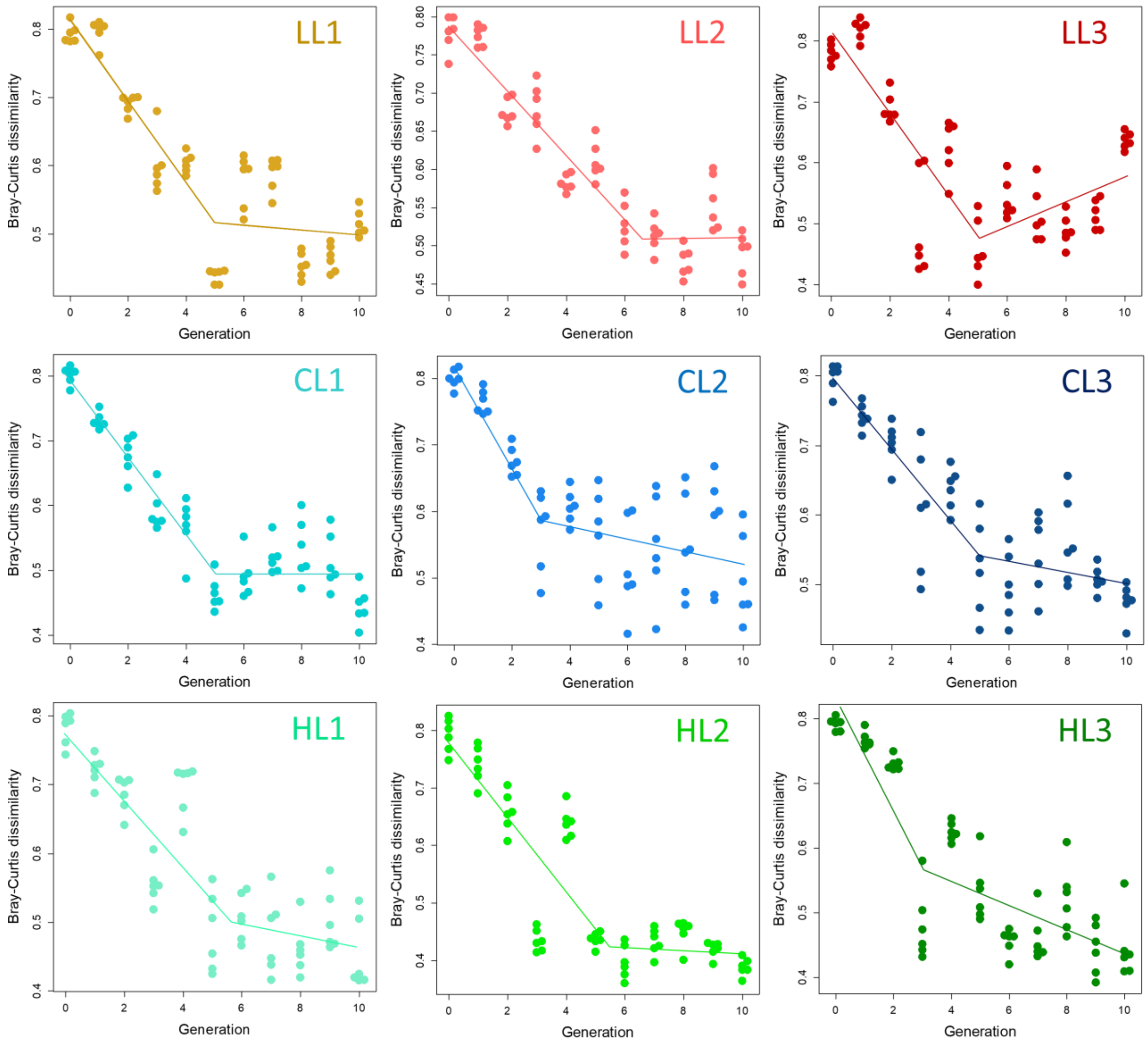


**Fig. S11:** Results of the unsupervised segmented analysis applied on the evolution of the beta diversity in each bacterial lineage. The analysis resulted in the systematic detection of a breaking point occurring, on average at generation = 4.90 (range: 3.00-6.65). After this breaking point, most lineages show a stabilization (slope not significantly different from zero), expect for the lineage LL3 which bounced back (a lineage were selection did not work, see Fig. S14 B), and HL3 which still has a decreasing trend but at a slower pace.

**Fig. S12.**

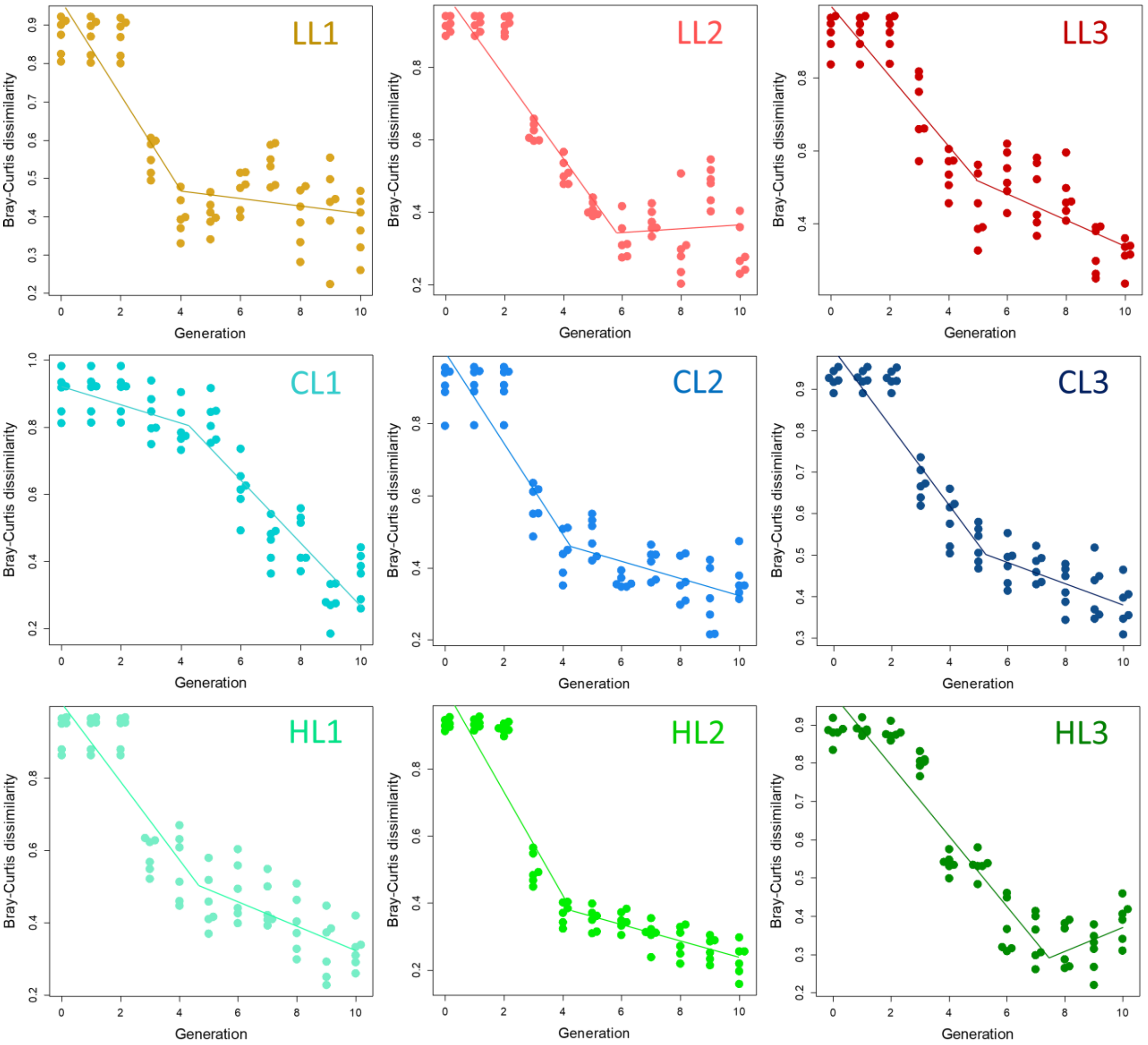


**Fig. S12:** Results of the unsupervised segmented analysis applied on the evolution of the beta diversity in each fungal lineage. The analysis resulted in the systematic detection of a breaking point occurring, on average at generation = 4.90 (range: 3.75-7.43). After this breaking point, most lineages display a slower slope than before, with only a few lineages reaching stabilization (slope not significantly different from zero: LL1, LL2 and HL3).

**Fig. S13.**

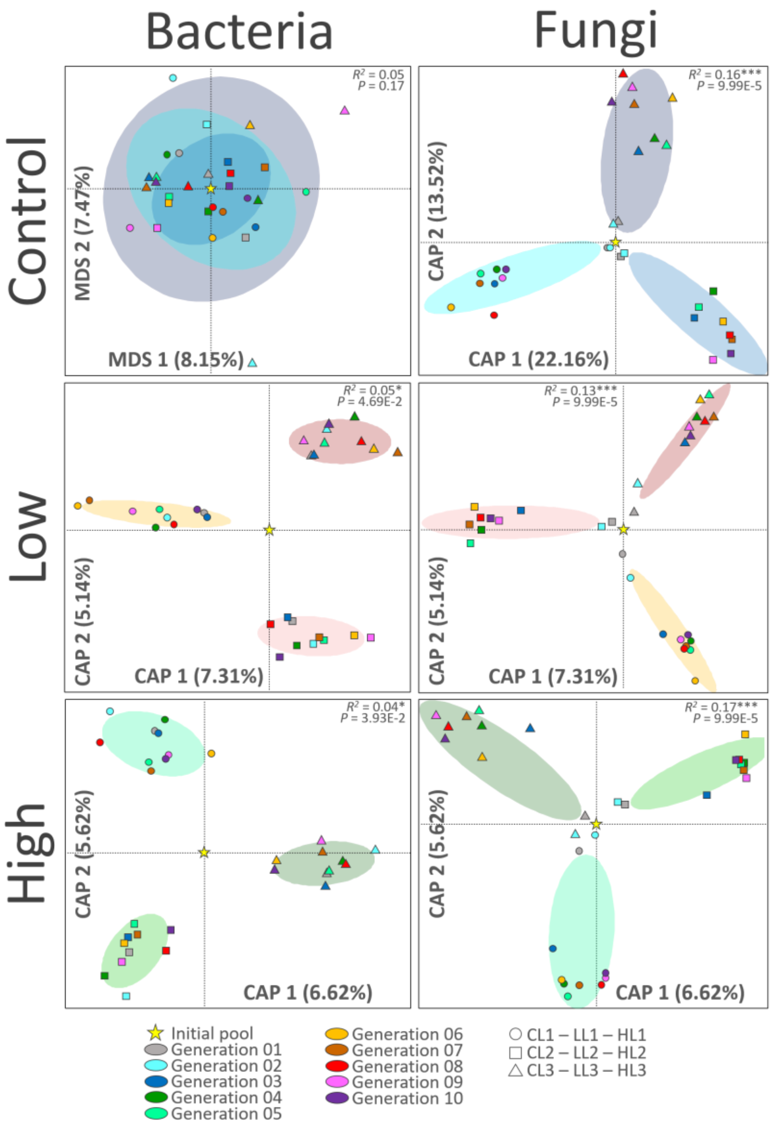


**Fig. S13:** Distance-based redundancy analysis of the selected microbiota rhizosphere for each lineage inside the selection groups. The models were built using the Bray-Curtis dissimilarity index, with 10.000 group permutations (Bray-Curtis ~ lineage). The $R^2$ values are indicating the percentage of variance explained by the model. If significant, the constrained coordinates are shown (model $P < 0.05$, CAP, Constrained Analysis of Principal coordinates). If not, the unsupervised coordinates are shown (model $P > 0.05$, MDS: Multi-Dimensional Scaling).

**Fig. S14.**

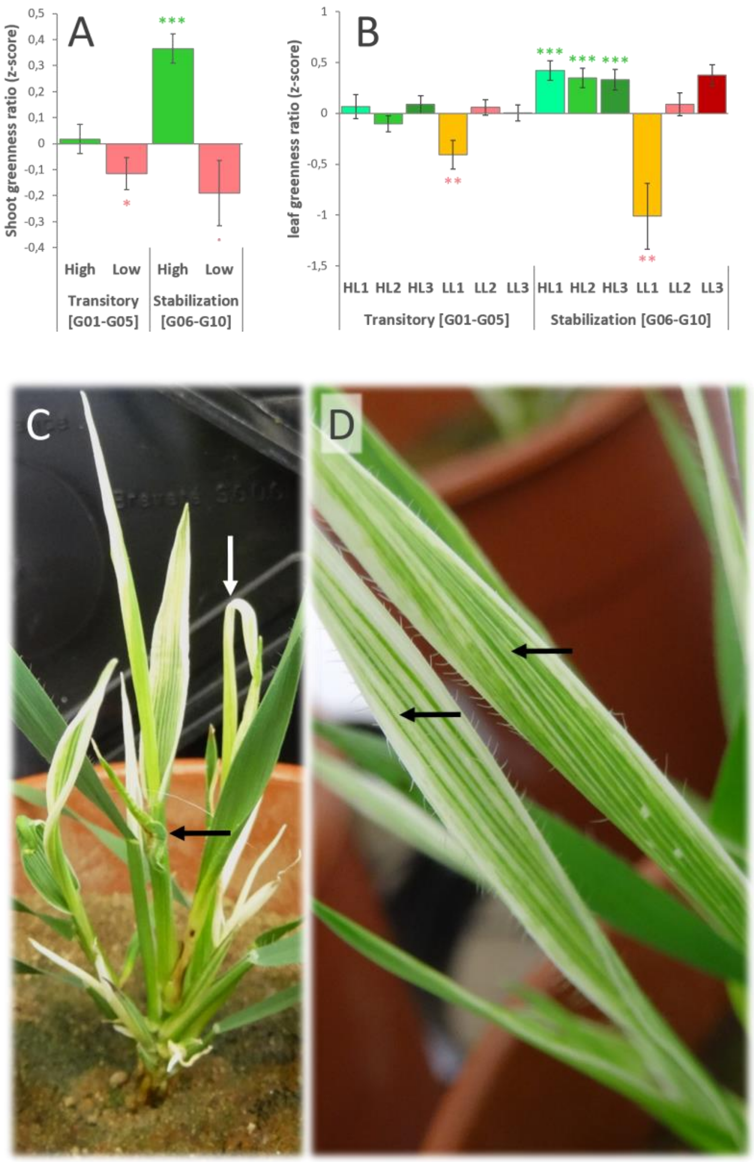


**Fig. S14:** Analysis of the leaf greenness index between the transitory and the stabilization phases. Panel A shows the results for the high and low selection groups, where the effect of high selection become very significant during the stabilization phase compared to the control baseline. Panel B shows the results for the high and low lineages, where the three high lineages showed a very significant and consistent increase compared to the control baseline. For the low selection group, a trend is detected, which was mostly explained by the very significant effects overserved in lineage LL1. Lineage LL2 was not significantly different than the control, while LL3 produced the opposite trend. Panels C and D are showing the very pronounced effects of the low selection in the plants in lineage LL1, featuring a delayed growth with apparition of twisted leaves (Panel C) and stripe discoloration on leaves (Panel D).

**Fig. S15.**

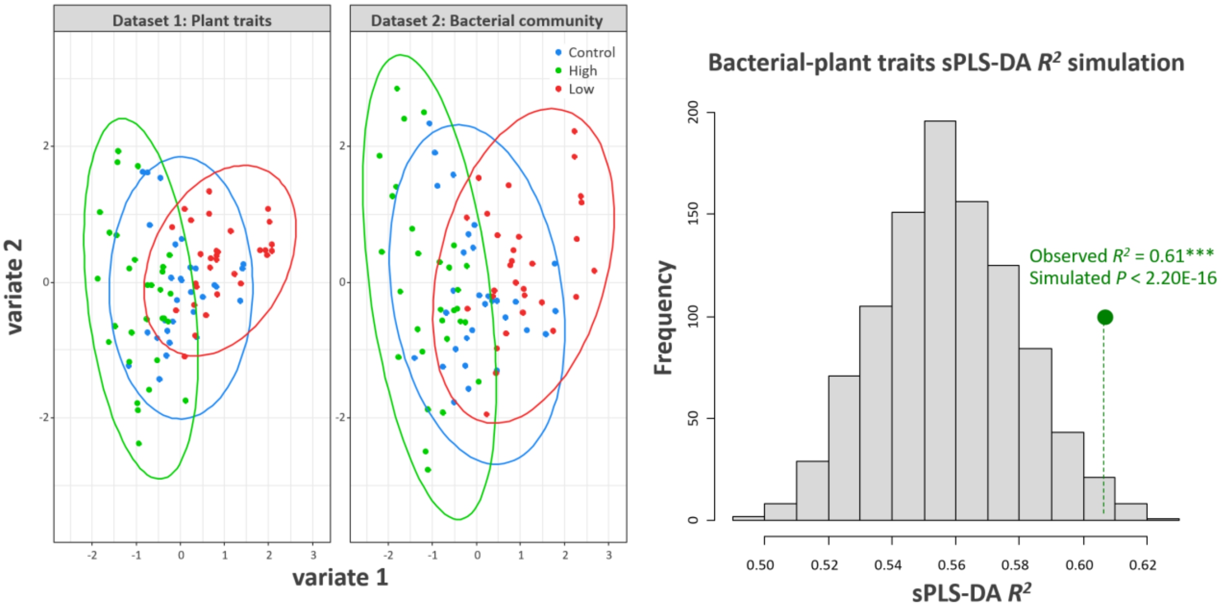


**Fig. S15:** Sparse partial least square discriminant analysis (sPLS-DA) between the multivariate plant trait dataset and the bacterial community. The figure shows the level of congruence between the two datasets for the first two components in their respective sub-space, and the random simulation applied to test the significance of the sPLS-DA $R^2$ (N = 1,000 group permutations, $R^2$ = 0.61***, $P$ < 0.001). $P$-value significance: « *** » $P$ < 0.001; « ** » $P$ < 0.01; « * » $P$ < 0.05; « . » $P$ < 0.1.

**Fig. S16.**

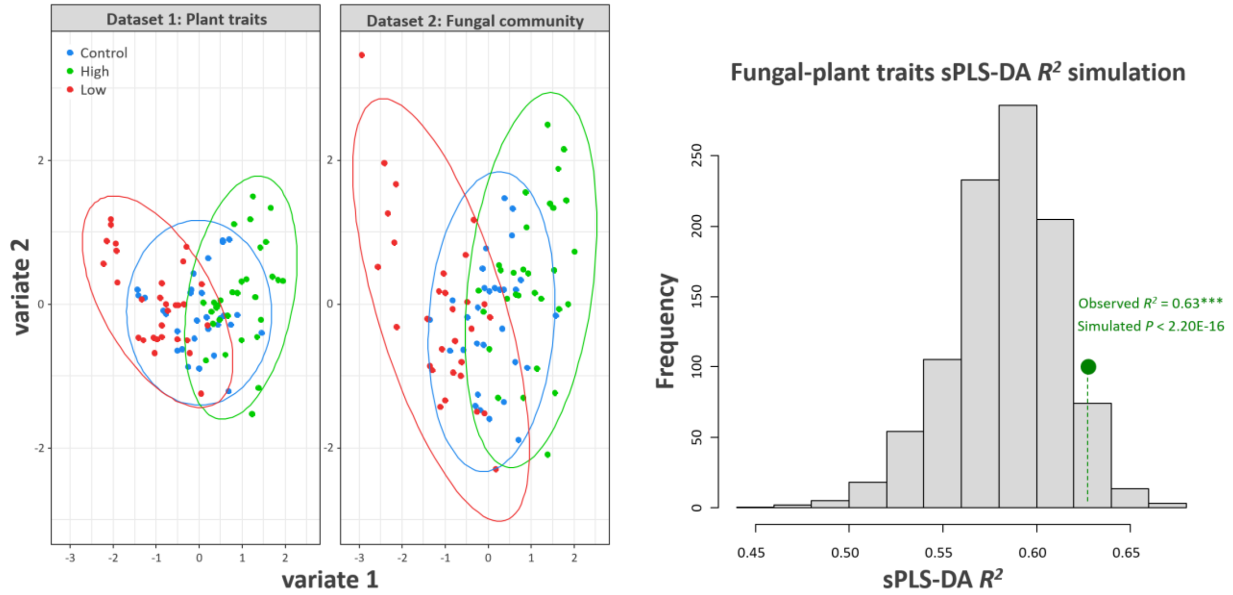


**Fig. S16:** Sparse partial least square discriminant analysis (sPLS-DA) between the multivariate plant trait dataset and the fungal community. The figure shows the level of congruence between the two datasets for the first two components in their respective sub-space, and the random simulation applied to test the significance of the sPLS-DA $R^2$ (N = 1,000 group permutations, $R^2$ = 0.63 ***, $P < 0.001$). $P$-value significance: « *** » $P < 0.001$; « ** » $P < 0.01$; « * » $P < 0.05$; « . » $P < 0.1$.

**Fig. S17.**

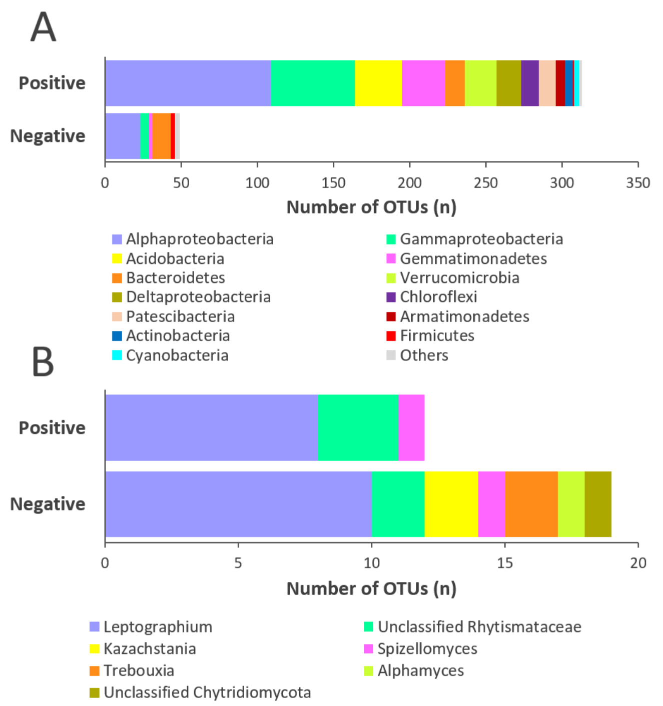


**Fig. S17:** Unweighted phylogenetic composition of the microbial taxa correlating either positively or negatively with leaf greenness across the entire dataset, as identified by the two sPLS-DA models. Panel A shows the phylogenetic origin of the 313 positive and 49 negative bacterial taxa. Panel B shows the phylogenetic origin of the 12 positive and 19 negative fungal taxa. Positive taxa featured very diverse and distinct bacterial taxa from Proteobacteria (Comamonadaceae and Caulobacteraceae, Deltaproteobacteria), Acidobacteria, Verrucomicrobia, Chloroflexi, Patescibacteria, Armatimonadetes and Actinobacteria. Negative taxa featured less diverse bacteria, from Firmicutes, Planctomycetes and Chlamydiae, as well as distinct fungal taxa from Ascomycota (*Kazachstania*), Chytridiomycota (*Alphamyces* and unclassified taxa) and Chlorophyta (*Trebouxia*). Both positive and negative categories had different taxa from common phylogenetic groups such as Alphaproteobacteria, Bacteroidetes Ascomycota (*Leptographium*, unclassified Rhytismataceae) and Chytridiomycota (*Spizellomyces*).

**Fig. S18.**

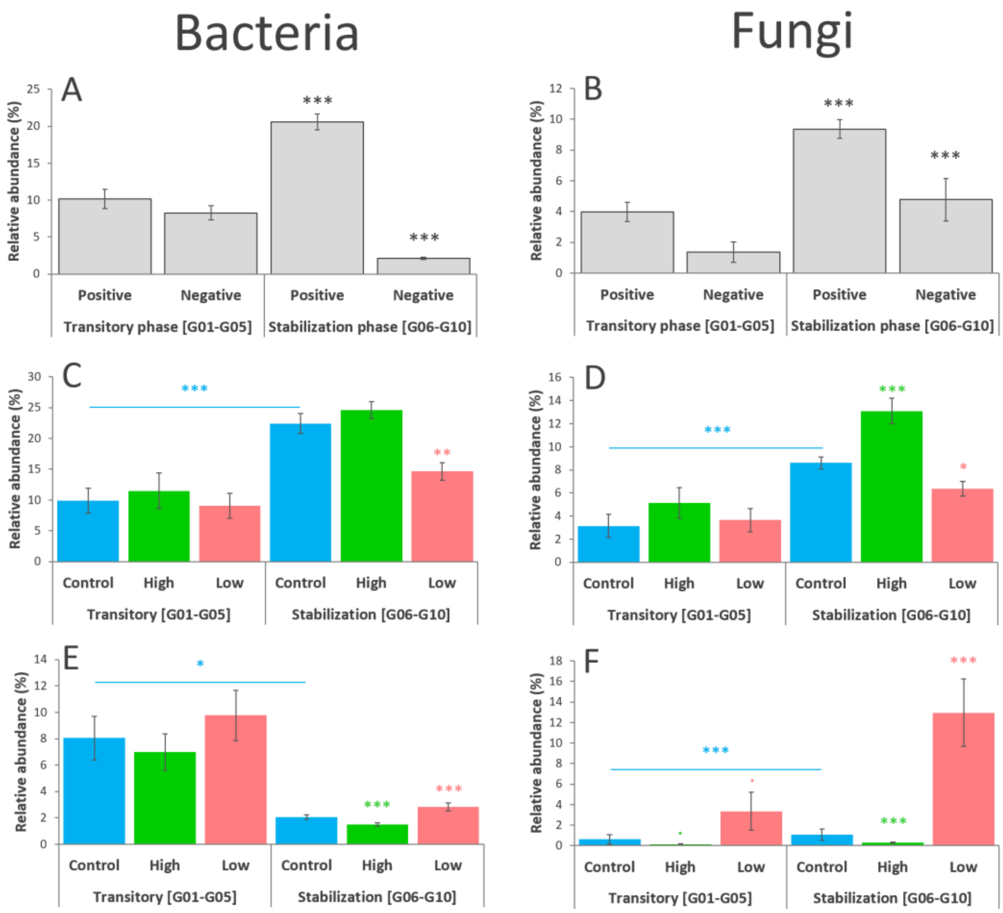


**Fig. S18:** Analysis of the grouped relative abundance of positive and negative taxa correlating with leaf greenness between the two phases. Panels A and B are showing the grouped abundance of positive and negative bacterial (A) and fungal taxa (B), regardless of the selection group. Panels C-D are showing the grouped abundance of positive bacterial (C) and fungal taxa (D) according to each selection group. Panels E-F are showing the grouped abundance of negative bacterial (E) and fungal taxa (F) according to each selection group. The statistical comparison of the grouped taxa abundance was done with a two-sample, two-sided Wilcoxon Rank Sum test. *P*-value significance: « *** » $P < 0.001$; « ** » $P < 0.01$; « * » $P < 0.05$; « . » $P < 0.1$. The horizontal blue lines and stars represent the comparison of the grouped microbial abundances observed in the control groups between the two phases. The green and red stars represent the statistical comparison of the high and low selection groups against their respective controls in each phase. Error bars are representing the standard error of the mean (N = 45 in panels A-B ; N = 15 in panels C-F).

**Fig. S19.**

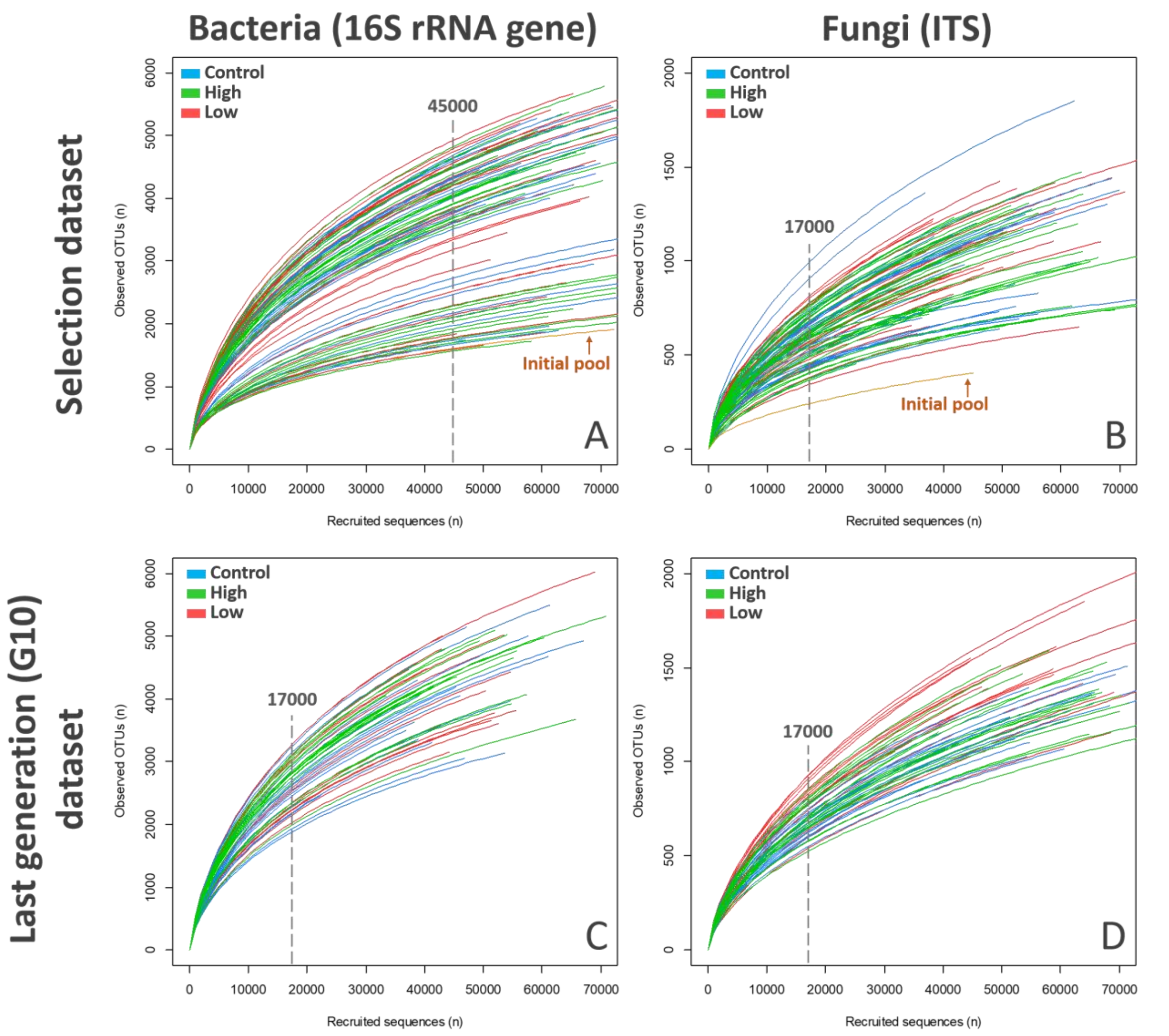


**Fig. S19: Rarefaction curves showing sample sequencing depth.** Profiles of bacterial 16S rRNA gene (A, C) and fungal ITS (B, D) amplicon sequencing for the inoculation pools from the selection experiment (A, B), and individual offspring rhizospheres from the generation 10 (C, D). The random resampling level applied to each dataset is indicated by the dotted lines. Initial microbial pools used to start the selection experiment are also indicated (A, B).

**Table S1.**

| **Factor** | **SOS** | **F** | **Var. (%)** | ***P*-value** | **Signif.** |
|---|---|---|---|---|---|
| Generation | 506761437 | 149.04 | 39.77 | < 2.00E-16 | *** |
| Generation:Selection | 22397911 | 2.96 | 1.76 | 1.18E-05 | *** |
| Generation:Selection:Lineage | 106885647 | 4.72 | 8.39 | < 2.00E-16 | *** |
| Residuals | 636044995 | - | 50.08 | - | - |

**Table S1: ANOVA model of the raw leaf greenness data.** Table shows the results of the ANOVA model (n = 1779 replicates, greenness~generation/selection/lineage). SOS = Sum Of Squares; MS = Mean Squares; F = Fisher index; Var.(%) = Percentage of variance explained based on MS; *P*-value significance: « *** » $P < 0.001$; « ** » $P < 0.01$; « * » $P < 0.05$; « . » $P < 0.1$.

**Table S2.**

| Dataset | Sample type | Marker | Number | Rarefaction level |
|---|---|---|---|---|
| All generations of the selection experiment | Inoculation pools | 16S - Bacteria | 91 | 45000 |
| | | ITS - Fungi | 91 | 17000 |
| Last generation (G10) of the selection experiment | Individual rhizospheres | 16S - Bacteria | 54 | 17000 |
| | | ITS - Fungi | 54 | 17000 |

**Table S2:** Summary of the sequencing datasets generated in this study (bacteria and fungi). The table presents the type and number of samples. The last column indicates the level of sequence rarefaction applied to normalize the datasets to even sequences numbers.
